# Spatial organization of supercoil dynamics during DNA replication

**DOI:** 10.1101/2024.10.14.618193

**Authors:** Yoshiharu Kusano, Soya Shinkai, Ryu-Suke Nozawa, Jiying Sun, Shuichi Onami, Satoshi Tashiro, Toru Hirota

## Abstract

Progression of DNA replication inevitably generates helical tension and resulting super-helical structures, or supercoils, that originate from the double helix of DNA. As DNA polymerases progress, negative supercoils accumulate in the wake of the replication fork, thereby impeding fork progression unless the supercoils are resolved topologically. Using super-resolution microscopy combined with spatial distribution analysis, here we describe that SMC5/6-mediated confinement of negative supercoils lies at the basis of releasing them by topoisomerase Top2A. Along with DNA replication, SMC5/6 progressively associates with chromatin and locally increases the density of supercoil clusters behind the fork, allowing Top2A to efficiently target and release the accumulated supercoils. These processes are essential to complete DNA replication prior to mitosis and therefore to ensure genome stability. Remarkably, we found that HeLa cells over-accumulate poorly confined negative supercoils beyond the processing capacity of cellular SMC5/6, which may exemplify the condition associated with genome instability of cancer cells.

## Introduction

Rapid and smooth DNA replication ensures duplication of entire genome before its segregation in the cell cycle. Disruption of this process results in a condition known as replication stress, which is associated with genome instability found in many types of human cancers (Burrell *et al*, 2013; Mankouri *et al*, 2013). Due to the helical nature of double-stranded DNA, every cell faces superhelical tension as elongating the replication fork and inevitably produces DNA supercoils, plectonemic coiled structures of DNA strands (Kurth *et al*, 2013; Ullsperger, 1995). Unwinding DNA by replicative helicases generates positive supercoils ahead of the forks, in the segment between converging forks, whereas DNA synthesis by DNA polymerases generates negative supercoils behind the forks (Dewar *et al*, 2015; Hingorani & O’Donnell, 2000; Kurth *et al*., 2013; Leu *et al*, 2003; Ullsperger, 1995). The efficient processing of these supercoils must be a prerequisite for the progression of replication (Kurth *et al*., 2013; Leu *et al*., 2003; Vologodskii *et al*, 1979). However, our understanding of how cells overcome these burdens is largely lacking, primarily because of the limitation of suitable methods to detect supercoil structures.

DNA topoisomerases represent a group of enzymes that resolve topological issues such as supercoils. They do so by cleaving and resealing either one (topoisomerase I, or Top1) or two (topoisomerase II, or Top2) strand(s) of the DNA molecule, in a direction that lowers the topological energy potential of the DNA substrate (Ullsperger, 1995; Wang, 2002). Both Top1 and Top2 in human cells can release superhelical tension and thereby relax positive and negative supercoils. A number of studies have demonstrated that inactivation of topoisomerases results in a significant inhibition of fork progression, reflecting the requirement of supercoil relaxation during replication (Bermejo *et al*, 2007; Brill *et al*, 1987; Goto & Wang, 1985).

The structure of supercoils is a determining factor in the ability of topoisomerase to access and subsequently act on them. For example, Top2 preferentially binds at sites of helix-helix juxtapositions (DNA crossovers) of supercoils (Zechiedrich & Osheroff, 1990), generated by locally increased superhelical density (Boles *et al*, 1990; van Loenhout *et al*, 2012; Vologodskii *et al*, 1992). The stability of supercoil is also a factor that affects the processibility of Top2 (Strick *et al*, 2000). In contrast to positive supercoils, that are confined and accumulated between two converging forks, negative supercoils are prone to diffuse dynamically along DNA behind the fork (van Loenhout *et al*., 2012). Therefore, the confinement and stabilization of negative supercoils during replication is crucial for Top2 to effectively resolve supercoils and facilitate smooth fork progression.

The SMC5/6 complex (hereafter, referred to as SMC5/6) is a member of the evolutionally conserved structural maintenance of chromosome (SMC) complexes, which play fundamental roles in organizing chromosome structure throughout the cell cycle (Aragon, 2018; Kim *et al*, 2023; Peng & Zhao, 2023; Pradhan *et al*, 2023; Taschner & Gruber, 2023). Although SMC5/6 interacts with and stabilizes the DNA substrates for topoisomerases in vitro, namely supercoils and catenations (Gutierrez-Escribano *et al*, 2020; Jeppsson *et al*, 2024; Kanno *et al*, 2015; Serrano *et al*, 2020), the biological significance of these activities of SMC5/6 remains largely unknown. Here, we show that, negative supercoils are confined by SMC5/6, thereby enabling Top2A to efficiently resolve them during DNA replication. Our work establishes a link between the functional relevance with molecular clustering, providing a novel framework in cell biology.

## Results

### Replication progression requires the association of SMC5/6 with replicated DNA

SMC5/6 has been implicated in dissolving DNA structures linking sister chromatids, and cells devoid of this mechanism exhibit frequent chromosome missegregation, manifested in severe anaphase bridges (**Fig. 1A**) (Gallego-Paez *et al*, 2014; Torres-Rosell *et al*, 2007; Venegas *et al*, 2020). To investigate cell cycle phase(s) that requires this SMC5/6 function to resolve the origin of chromosomal bridges, we generated HeLa and RPE-1 cell lines carrying the auxin-inducible degron (AID) biallelically on endogenous SMC6 genes, SMC6-degron cells (**Fig. EV1A-C**). By acutely degrading SMC6 at various timepoints before mitosis (**Fig. 1B**), we found that cells that experienced S phase progression without SMC5/6 gave rise to persistent unresolved DNA structure between sister chromatids in both cell lines (**Fig. 1C and EV1D**).

**Figure 1.**
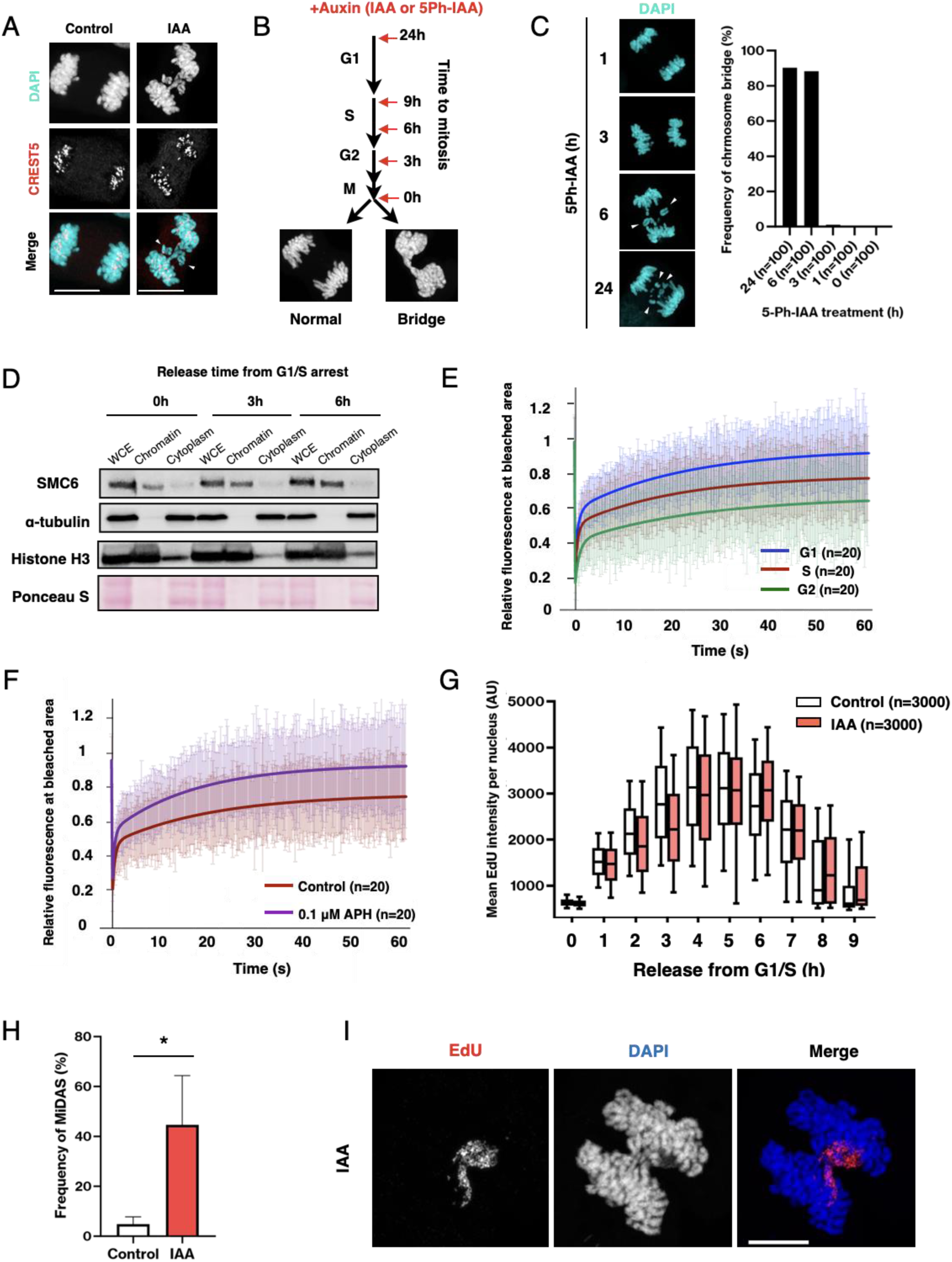
Cumulative association of SMC5/6 with chromatin during replication progression. (A) Representative images of anaphase bridges in HeLa cells depleted of SMC6. Arrowheads indicate chromatid pairs isolated by chromatid gaps (see Figure S2A-S2D). Scale bars, 10 μm. (B) Logarithmically grown RPE-1 or HeLa cells were treated with 5Ph-IAA or IAA to deplete SMC6 for the indicated period of time. Mitotic cells were analyzed for anaphase bridges as shown in the histogram, respectively (RPE, Figure 1C; HeLa, Figure S1D). SMC6 was depleted at: G2 (for 3 h treatment), late S (6 h), early S (9 h), and G1 (24 h) phases. (C) Representative anaphase cells of SMC6-degron RPE-1 cells that had been treated with 5Ph-IAA for indicated period of time (left). Arrowheads indicate isolated chromatid pairs associated with symmetrical gaps (see Figure S2A-S2D). Bar charts show the frequency of anaphase bridges in each sample (right). (D) Chromatin fractionation shows increasing amounts of stably chromatin-bound SMC6 during S phase progression. SMC6-degron HeLa cells synchronized at G1/S boundary were released for the indicated period of times. Enrichment of the chromatin and the cytoplasmic fractions is verified by histone H3 and α-tubulin, respectively. (E, F) FRAP analysis of SMC6-mClover. (E) Kinetics of fluorescence recovery in G1 (blue), S (red), and G2 (green) phases are shown. (F) Inhibition of DNA polymerases by aphidicolin (APH) treatment significantly reduces the stably bound-SMC5/6 fraction in mid S phase (purple) from the control level (brown). Error bars are standard deviations (SD) for each timepoint. (G) Replication kinetics in the first round of S phase in the absence of SMC5/6. HeLa cells arrested at the G1/S boundary were released for the indicated time and EdU fluorescence intensities in individual cells were measured for each timepoint and shown in box plots (the central lines are the means). Control (white); IAA (red). AU, arbitrary unit. (H, I) Mitotic DNA synthesis (MiDAS) in the absence of SMC5/6. (H) Bar charts show the frequency of mitotic chromosomes positive for EdU (mean ± SD, n=3). * indicates *P* < 0.05, two-tailed Wilcoxon-Mann-Whitney test. (I) A representative image of the EdU-positive chromosome bridge. Scale bar, 10 μm.

To validate the chromosome interaction of SMC5/6 in S phase, we examined the abundance of SMC5/6 stably bound to chromatin. Nuclear SMC5/6 was found to increasingly associate with chromatin during S phase (**Fig. 1D**). The fluorescence recovery after photobleaching (FRAP) experiments also indicated that stably associating fraction emerges and progressively increases as replication proceeds (**Fig. 1E and EV1E**). This stable association of SMC5/6 was strongly inhibited by treating cells with aphidicolin (APH), a specific inhibitor of DNA polymerase α, δ, and ε (**Fig. 1F**). This suggested that SMC5/6 interact with replicated DNA situated behind the replication forks, as APH blocks the activity of DNA polymerases without affecting that of helicases (Walter & Newport, 2000). Thus, SMC5/6 seemed to act on replicated DNA during S phase as suggested in yeast experiments (Betts Lindroos *et al*, 2006; Jeppsson *et al*, 2014; Kegel *et al*, 2011).

These results prompted us to examine if the chromatin association of SMC5/6 affects replication kinetics. Although the bulk replication in the absence of SMC5/6 overall proceeded normally (**Fig. EV1F**) (Venegas *et al*., 2020), a measurable replication delay was observed at the single-cell level, following a pulse incorporation of the DNA analog 5-ethynyl-2’-deoxyuridine (EdU) into nascent DNA (**Fig. 1G**). In SMC6-depleted cells, replication origin firing seemed intact ∼1 h after the release from G1/S boundary; however, in the following timeframes (2 ∼ 10 h), not only the time initiating replication of late-replicating domains but also the time completing replication were consistently delayed (**Fig. 1G and EV1G**). In live cells, in which fluorescently tagged proliferative cell nuclear antigen (PCNA) is expressed to differentiate early and late replication, we found that SMC6-depleted cells required longer times to replicate both early- and late-replicating domains (**Fig. EV1H, I**).

Thus, while replication in early domains can initiate normally, cells devoid of SMC5/6 fail to efficiently process the replication, for both early and late domains. These results challenge the prevailing model that SMC5/6 primarily facilitates replication termination through fork rotation, without contributing to fork progression prior to termination (Kegel *et al*., 2011).

The extent of the replication delay was beyond the detectable level in bulk population analysis (**Fig. EV1F**), however it became conspicuous for its persisting replication in mitosis. This phenomenon called mitotic DNA synthesis, or MiDAS (Minocherhomji *et al*, 2015) were found in a majority of mitotic cells (**Fig. 1H**) (Atkins *et al*, 2020). Interestingly, MiDAS was typically found adjacent to chromosome bridges in anaphase (**Fig. 1I**), implying that the presence of unreplicated thereby unresolved chromosome regions were prominent sources of chromosome bridges.

### Uncompleted replication causes chromatid gap-related chromosome bridges

In SMC6-depleted metaphase chromosomes, we noticed numerous symmetric gaps in chromatid arms (**Fig. EV2A**) (Venegas *et al*., 2020). These gaps detached chromatid pairs, which share similar structural characteristics with small chromatid fragments lagged behind the separating sisters in anaphase (**Fig. EV2B**). In live-cell imaging analysis, we could indeed see lagging of isolated chromatid pairs that were often left behind the separating sisters in anaphase, presumably because the poleward pulling force failed to apply to the detached chromatids across those intervening gaps (**Fig. EV2C, D**).

We could reason that these chromatid gaps originate from uncondensed chromosomal domain relating to as-yet-unreplicated chromatin, provided that it remains refractory to mitotic chromosome condensation when replication is on-going (Johnson & Rao, 1970). Spectral karyotyping of chromosome spreads, which specifically determined on which chromosomes the gaps had taken place, showed that the number of gaps were highly correlated with the length of chromosome in the absence of SMC6 (**Fig. EV2E, F**), similar to the finding in budding yeast (Kegel *et al*., 2011). These results support the idea that SMC5/6 facilitates prompt replication and thereby prevents sister chromatid non-disjunction in mitosis.

### SMC5/6 is required to prevent accumulation of DNA supercoils in S phase

To investigate how SMC5/6 facilitates replication progression, we examined the amount of supercoiled DNA structure, in light of SMC5/6’s activity to stabilize supercoils in vitro (Gutierrez-Escribano *et al*., 2020; Jeppsson *et al*., 2024; Serrano *et al*., 2020; Taschner & Gruber, 2023). We used psoralen (Bermudez *et al*, 2010; Naughton *et al*, 2013), a chemical that intercalates into negative supercoils (**Fig. EV3A**) that arise behind the forks (Hingorani & O’Donnell, 2000; Kurth *et al*., 2013; Leu *et al*., 2003; Ullsperger, 1995), where SMC5/6 stably interacts (**Fig. 1E, F**) (Betts Lindroos *et al*., 2006; Jeppsson *et al*., 2014; Kegel *et al*., 2011). The amount of negative supercoils was undetectable in SMC5/6 proficient cells, whereas there was significant psoralen incorporation in SMC6-depleted cells, especially when they were enriched in S phase (**Fig. 2A and EV3B, C**).

**Figure 2.**
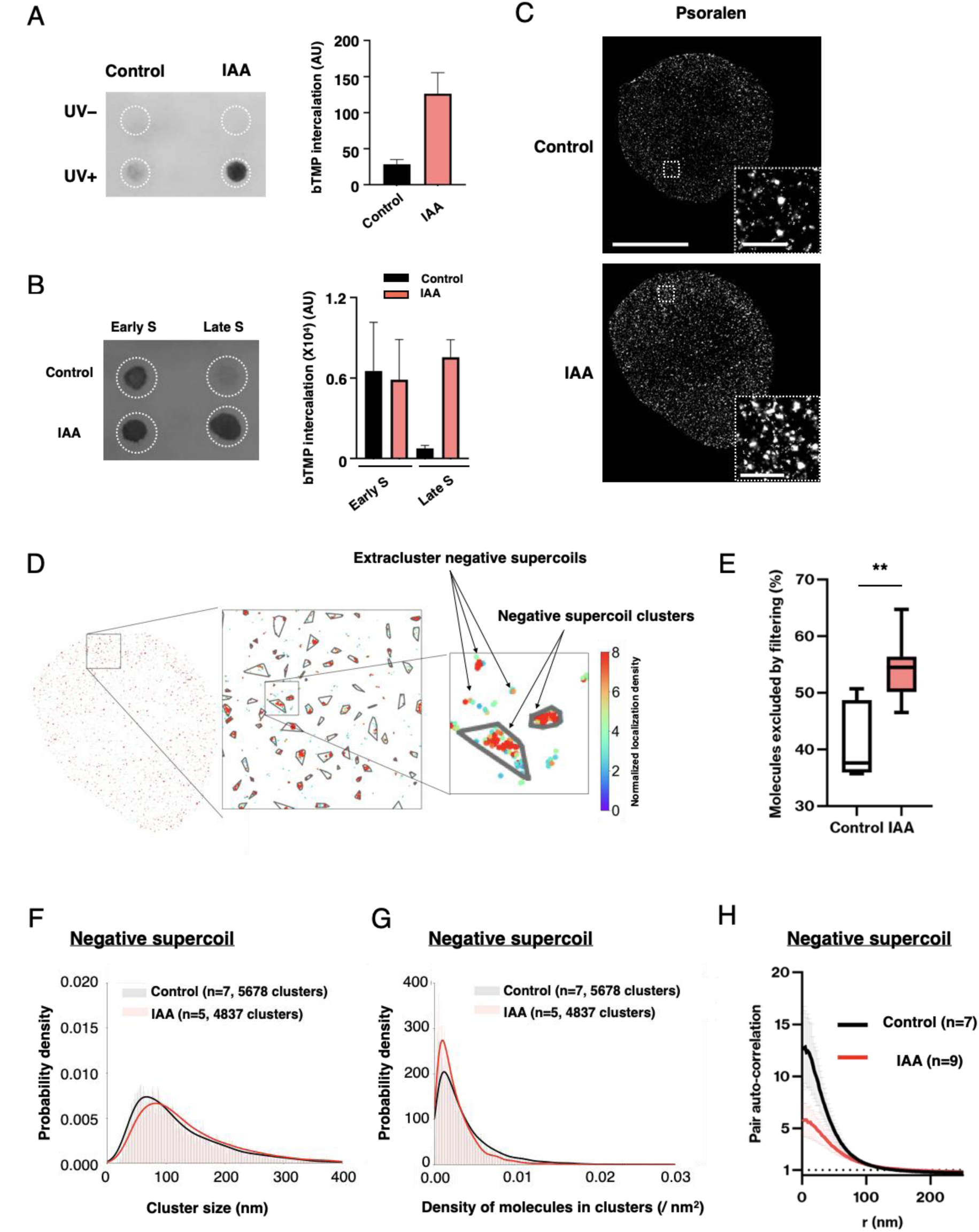
SMC5/6 is required to confine and release negative supercoils during S phase. (A) HeLa cells synchronized at late S phase were treated with biotinylated psoralen (bTMP) and intercalated bTMP was visualized with streptavidin-HRP on extracted and spotted DNA. Note that bTMP was detected when fixed by UV irradiation (UV+) but not when not fixed (UV-). Left, representative dot blot image. White dotted circles indicate where DNA extracts were spotted. Right, bar charts show signal intensities in arbitrary units (mean ± SD, n=3). (B) Psoralen intercalation was examined in DNA extracts prepared from cells synchronized in early and late S phase. S phase progression in the presence of low-dose CDC7 inhibitor allowed to obtain cell populations enriched in early and late S phases (see Figure S3D). (C) Representative STORM images of biotinylated psoralen intercalated in SMC6-degron HeLa cells. Cells in late S phase were examined by fluorescence microscopy for mCherry-labeled PCNA and by STORM imaging for psoralen molecules, detected with azide-fused Alexa-Fluor-647. Scale bar, 10 μm. Enlarged views show that psoralen molecules are densely distributed in the SMC6-depleted cell nucleus. Scale bar, 1 μm. (D) Representative segmented image of late S phase nuclei of IAA-treated cells. In the enlarged views, a plethora of identified clusters and molecules excluded from the clusters are identified. The extent of molecular assemblies is indicated by the color-coded density. (E-G) Quantitative analysis of psoralen clusters. Segmentation analysis identified ∼1×10^3^ psoralen clusters per cell in control and IAA treated HeLa cells in late S phase. The proportions of psoralen molecules excluded from the clusters are shown in box plots (E). ** indicates *P* < 0.01; two-tailed Wilcoxon-Mann-Whitney test. Diagrams show the probability density of the cluster size (F) and the cluster density (G). The fitted mean curves for control and IAA-treated cells are shown in black and red, respectively. Two-tailed Wilcoxon-Mann-Whitney tests indicate differences in the size (*P* < 0.0001) and density (*P* < 0.0001) between control and IAA. (H) Quantitative analysis of psoralen molecular assemblies in HeLa cells. Datasets of the distance between each psoralen molecule to all the other psoralen molecules are used for the analysis. The auto-correlation function *g*(r) = 1 indicates random distribution and greater than 1 indicates biased distribution of the molecules. The densities of psoralen assemblies in late S phase are analyzed in SMC6 depleted (IAA, red) and control (black) cells.

To investigate whether these negative supercoils accumulate during S phase progression, we prepared cell populations enriched in either early or late S phase by controlling the firing of late-replicating domains with a CDC7 inhibitor (Bousset & Diffley, 1998) (**Fig. EV3D**). While the negative supercoils were accumulated in early S phase, they were normally dissolved by late S phase. By contrast, in SMC6-depleted cells, they were progressively accumulated through late S phase (**Fig. 2B**). Notably, the use of psoralen in live cells repressed EdU incorporation (**Fig. EV3E**), verifying the idea that the persistence of negative supercoils perturbs replication progression. These results indicate that the role of SMC5/6 in stabilizing supercoils (Gutierrez-Escribano *et al*., 2020; Serrano *et al*., 2020) is indeed required for releasing supercoils during replication, thereby promoting fork progression.

### SMC5/6 confines negative supercoils and increases local superhelical density

To address how SMC5/6 might release negative supercoils during replication, we used psoralen to probe negative supercoils and studied their dynamics in S phase cells (**Fig. EV3B**). We employed stochastic optical reconstruction microscopy (STORM), a super-resolution imaging technique that allows for localizing single molecules within the nucleus. We found that psoralen-probed negative supercoils assemble and form diffraction-limited clusters in late S phase, and that negative supercoils become more densely present after SMC6 depletion (**Fig. 2C and EV3F**).

Segmentation analysis allowed us to define the clusters based on the biased rather than random molecular distributions and, therefore, to calculate the proportion of molecules that were excluded from the clusters (**Fig. 2D**). Since the proportion of molecules excluded from clusters was markedly elevated in the absence of SMC5/6 (**Fig. 2E and EV3G**), we reasoned that, without SMC5/6, negative supercoils are less confined and cluster with less efficiency. Consistent with this possibility, we found that SMC6 depletion led to a notable increase in the size and a reduction in the density of the negative supercoil clusters (**Fig. 2F, G and EV3H, I**). Pair auto-correlation analysis showed a reduction in the density of the clusters throughout the S phase nucleus (**Fig. 2H and EV3J**). Since supercoils can dynamically move along the helical path in vitro (a process referred to as supercoil diffusion) (van Loenhout *et al*., 2012), these alterations in the spatial organization of negative supercoils in the absence of SMC5/6 imply that SMC5/6 constrains supercoil diffusion behind the fork, presumably through its physical association with supercoils (Gutierrez-Escribano *et al*., 2020; Serrano *et al*., 2020).

### SMC5/6 promotes clustering of Top2A on supercoils during replication

Supercoils impede the fork progression unless they are released by topoisomerases (Bermejo *et al*., 2007; Ullsperger, 1995; Wang, 2002), and SMC5/6 is unlikely to release supercoils independently (Gutierrez-Escribano *et al*., 2020; Jeppsson *et al*., 2024; Serrano *et al*., 2020). Thus, a corollary of these observations was that SMC5/6 is involved in topoisomerase-mediated supercoil relaxation during replication. Along this hypothesis, we sought for cellular outcomes that were shared between cells deleted of SMC6 and of topoisomerases. First, we assessed the impact of SMC6 depletion on replication in the second S phase. Top2 depletion significantly disrupts bulk replication in the second cell cycle, while depletion of Top2 or Top1 does not affect bulk replication in the first round of the cell cycle (Baxter & Diffley, 2008; Bermejo *et al*., 2007). Similarly, we found a remarkable delay in the second replication in SMC6-depleted cells (**Fig. EV4A, B**). Second, we found that replication kinetics with the Top2 inhibitor ICRF-193 were reminiscent of those with SMC6 depletion, whereas treatment with the Top1 inhibitor camptothecin (CPT) strongly suppressed EdU incorporation throughout the replication process (**Fig. EV4C**). Third, SMC6 depletion caused chromosomal rearrangements, including characteristic chromosome fusions and diplochromosomes (**Fig. EV4D-G**). All of these aberrant chromosomes were also induced by transient ICRF-193 treatment (Sumner, 1998). These observations suggest that Top2 rather than Top1 is the primary topoisomerase that resolves replication-associated supercoils with the aid of SMC5/6.

To address the potential relationship between SMC5/6 and Top2-mediated supercoil relaxation during replication, we tagged endogenous Top2A with mCherry homozygously in the SMC6-degron HeLa cell line and examined its dynamic interaction with chromatin (**Fig. EV5A**), as Top2A is the isoform that ubiquitously expressed in proliferative cells (Wang, 2002). The kinetics of fluorescence recovery of Top2A-mCherry in S phase were similar to those previously documented (Christensen *et al*, 2002), and became slightly slower in the absence of SMC5/6 (**Fig. EV5B**). Fitting with bi-exponential functions indicated two different interaction modes, i.e., stably and dynamically chromatin bound fractions. Notably, the residence time of the stably bound fraction was prolonged in the absence of SMC5/6, whereas that of the dynamically bound fraction remained largely unaffected (**Fig. EV5C**). As the catalytic processing of Top2A depends on the stability of supercoils (Strick *et al*., 2000), the slower turnover rate may indicate that supercoils became labile and Top2A spent more time in the catalytic process in the absence of SMC5/6. These results seem to be consistent with our findings that loss of SMC5/6 destabilizes supercoil dynamics in S phase (**Fig. 2E-H and EV3G-J**).

With the use of STORM imaging analysis, the nuclear signal of Top2A in fluorescence microscopy turned out to be the sum of numerous diffraction-limited assemblies in late S phase (**Fig. 3A, B**). Segmentation analysis indicated that, in the absence of SMC5/6, the size of Top2A clusters became larger, their density smaller, and, accordingly, the fraction excluded from the clusters inversely increased (**Fig. 3C-E and EV5D-F**). Pair-correlation analysis corroborated the finding that Top2A assemblies became less dense throughout the nucleus in the absence of SMC5/6 (**Fig. 3F and EV5G**). As these changes were analogous to those observed in negative supercoil clusters in the absence of SMC5/6, we postulated that Top2A clusters correlate with negative supercoil clusters.

**Figure 3.**
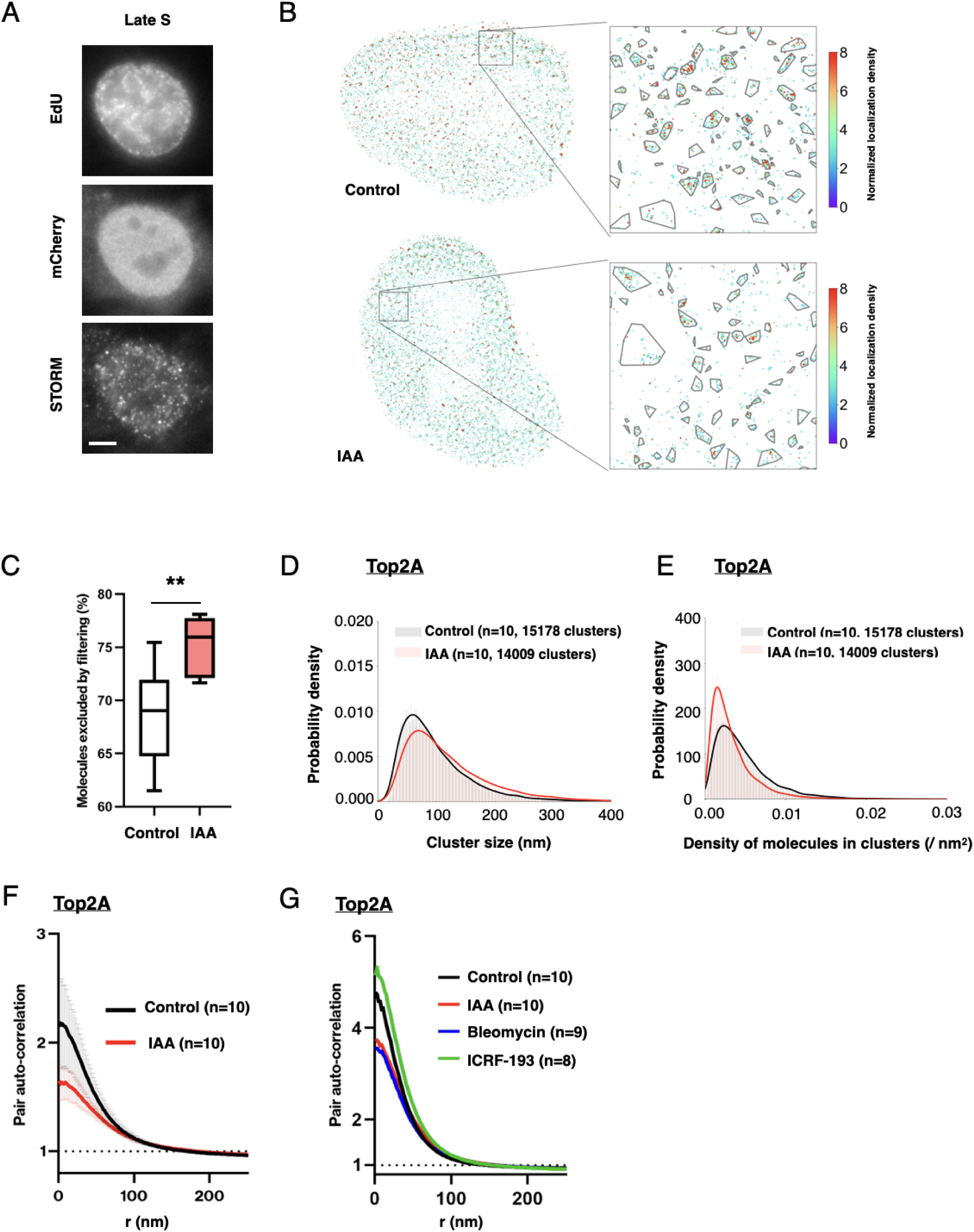
SMC5/6 promotes Top2A clustering onto supercoils in S phase. (A) Representative STORM images of mCherry-labeled Top2A in SMC6-degron HeLa cells. Cells in early and late S phases were examined with fluorescence microscopy for EdU incorporation, probed with 488 picolyl-azaide, and Top2A-mCherry, and with STORM imaging for Top2A molecules, detected with anti-Cy5. Scale bar, 3 μm. (B) Segmentation of late S phase nuclei identified ∼1.5×10^3^ Top2A clusters per cell and in control and IAA treated HeLa cells. Enlarged views show a plethora of identified clusters and the Top2A molecules excluded from the clusters. The extent of molecule assemblies is indicated by color-coded density. (C-E) Quantitative analysis of Top2A clusters. The proportions of Top2A molecules excluded from the clusters are shown in box plots (C). ** indicates *P* < 0.01; Two-tailed Wilcoxon-Mann-Whitney test. Diagrams show the probability density of the cluster size (D) and the cluster density (E). The fitted mean curves for control and IAA-treated cells are shown in black and red, respectively. Two-tailed Wilcoxon-Mann-Whitney tests indicate differences in the size (*P* < 0.0001) and density (*P* < 0.0001) between control and IAA. (F) Quantitative analysis of the Top2A molecular assemblies in HeLa cells in early and late S phases. Datasets of the distance between each Top2A molecule to all the other Top2A molecules are used for the analysis. (G) Analysis of the Top2A molecular assemblies in late S phase of HeLa cells, after 30 min treatment with DMSO (control, black), 35 μM ICRF-193 (green), 100 μM bleomycin (blue), or after 12 h treatment with IAA (red). Error bars represent standard deviations. Similar results were obtained in two independent experiments.

To investigate the relevance of these molecular densities of clusters, cells were treated with ICRF-193 to stop the release of supercoils and catenations by forming Top2A clamps. This resulted in an increased density of Top2A in the clusters (**Fig. 3G and EV5H**), suggesting that density is indicative of the abundance of supercoils and catenations. Conversely, treatment of cells with bleomycin to dissolve supercoils beyond detectable levels (**Fig. EV3A**) (Naughton *et al*., 2013) caused a notable reduction in Top2A density (**Fig. 3G and EV5H**). Provided that bleomycin unwinds supercoils by introducing single-strand breaks (nicks), without breaking the double strand of DNA, bleomycin treatment would not, in principle, release catenations, favoring the idea that Top2A clusters are formed relating supercoils. It is worth noting that SMC6 depletion caused a reduction in Top2A density, to an equivalent level to that in bleomycin treatment (**Fig. 3G**). These findings suggest that, in the absence of SMC5/6, Top2A fails to form dense clusters, even in the presence of supercoils (**Fig. 2A, B**). Collectively, these results highlight the crucial role of SMC5/6 in facilitating Top2A’s recognition and relaxation of supercoils during DNA replication.

### Spatial distributions of SMC5/6 and Top2A clusters relative to replication forks

To characterize the spatial distribution of Top2A clusters with respect to replication forks, we conducted a coordinate-based colocalization (CBC) analysis using two-color STORM imaging datasets (Malkusch *et al*, 2012; Pageon *et al*, 2016) (**Fig. 4A**). The CBC analysis of late S phase nuclei showed that distribution of PCNA is highly correlated with that of both Top2A and Top1 molecules (**Fig. 4B and EV6A**). These results are in agreement with previous ChIP analysis showing that topoisomerases are recruited immediately onto replication-ongoing regions (Bermejo *et al*., 2007). It is noteworthy that the distribution of SMC5/6 was also found to be significantly correlated with that of PCNA (**Fig. 4C**).

**Figure 4.**
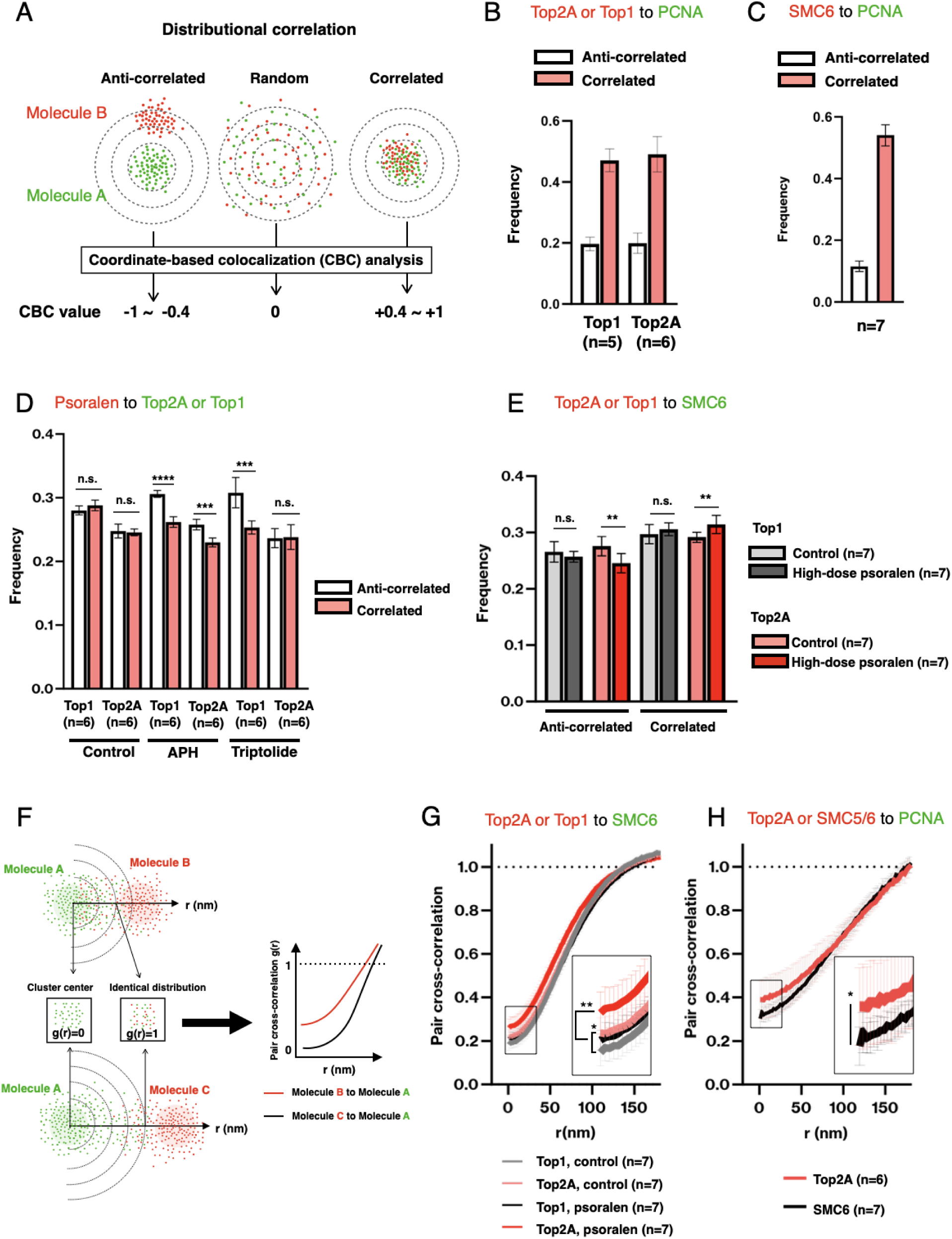
Distributions of SMC5/6 and Top2A clusters localized in the vicinity of the replication fork. (A) Schematic illustrating coordinate-based colocalization (CBC) analysis. When molecule A (green) and molecule B (red) are similarly distributed, the CBC value becomes positive (CBC value of +0.4 ∼ +1 is defined as correlated distribution). When two molecules are exclusively distributed, the CBC value becomes negative (CBC value of −0.4 ∼ −1 is defined as anti-correlated distribution). (B) CBC analysis shows that Top2A and Top1 are highly correlated with PCNA in late S phase. Anti-correlated distribution, white bars; correlated distribution, red bars. (C) CBC analysis shows that SMC5/6 is highly correlated with PCNA in late S phase. Anti-correlated distribution, white bars; correlated distribution, red bars. (D) CBC analysis shows the distribution correlation between Top2A or Top1 and psoralen in S phase cells. Cells were synchronized in S phase as shown in Figure S2B. Anti-correlated distribution, white bars; correlated distribution, red bars; APH, 1 μM APH treatment for 1 h; Triptolide, 1 μM Triptolide for 1 h (Bensaude, 2011). n.s., not significant; *** indicates *P* < 0.001, ** indicates *P* < 0.01; * indicates *P* < 0.05; two-tailed Wilcoxon-Mann-Whitney tests. (E) CBC analysis shows the distribution correlation between Top2A or Top1 and SMC6. When supercoils are immobilized, the correlated relationship between Top2A and SMC5/6 is emphasized compared to the counterpart. Top1 (control), gray bars; Top1 (high-dose psoralen), dark gray bars; Top2A (untreated), red bars; Top2A (high-dose psoralen), dark red bars. n.s., not significant; ** indicates *P* < 0.01; two-tailed Wilcoxon-Mann-Whitney tests. (F) A scheme of pair cross-correlation analysis. Datasets of the distance between each of molecule A (green) to all of molecule B or C (red) are used for the analysis. This cross-correlation analysis estimates the relative distribution of molecule B or C from molecule A (cross-correlation) to the distribution of molecule A itself (auto-correlation) (see Methods for details). Pair cross-correlation function reaches 1 when molecule B or C is distributed similarly to molecule A. (G) Correlative distributions of topoisomerases to SMC5/6. The pair cross-correlation analysis indicates that both Top2A or Top1 molecules are distributed less frequently than SMC5/6 molecules themselves, where *g*(r) is smaller than 1. The frequency of Top2A is higher in this range compared to Top1, suggesting the propensity of Top2A distribution being closer to that of SMC5/6. This frequency is further increased in response to psoralen-mediated supercoil immobilization. Top1, gray; Top1 + psoralen, black; Top2A, pink; Top2A + psoralen, red. Similar results were obtained in two independent experiments. **, *P* < 0.01 (red vs black); *, *P* < 0.05 (pink vs gray); two-tailed Wilcoxon-Mann-Whitney test was used to compare data points of g(r=0). (H) Correlative distributions of Top2A and SMC5/6 with PCNA, respectively. Top2A, red; SMC5/6, black. *, *P* < 0.05; two-tailed Wilcoxon-Mann-Whitney tests was used to compare data points of g(r=0).

Negative supercoils can arise not only behind the fork progression but also behind the transcription machinery (Hingorani & O’Donnell, 2000; Kurth *et al*., 2013; Leu *et al*., 2003; Liu & Wang, 1987; Ullsperger, 1995). To estimate the proportion of topoisomerase clusters that are associated with replication-related negative supercoils, we asked how these correlative distributions might be affected when replication or transcription is inhibited. Both of these inhibitors resulted in a reduction in the correlated distributions, and an increase in the anti-correlated distributions, between Top1 and negative supercoils. Conversely, inhibition of replication, but not of transcription, led to a reduction in the correlated distribution between Top2A and negative supercoils (**Fig. 4D**). These findings suggest that Top2A plays a specific role in processing the negative supercoils that arise behind the replication forks.

Given the correlation between Top2A clustering and its functional relevance, we sought to investigate the spatial relationship between SMC5/6 and Top2A. CBC analysis revealed that the immobilization and accumulation of supercoils by high-dose psoralen treatment resulted in an increased correlated distribution between Top2A and SMC6 and a decreased anti-correlated distribution. By contrast, no notable change was observed between Top1 and SMC6 (**Fig. 4E**). To determine which topoisomerase clusters are situated in closer proximity to SMC5/6 clusters, we performed pair cross-correlation analysis, which provided a relative distance between distributions of Top2A or Top1 and SMC5/6 (**Fig. 4F and EV6B**). The Top2A and Top1 clusters were exclusively enriched from that of SMC5/6 clusters. However, the Top2A clusters revealed a higher tendency to position in closer proximity to the SMC5/6 cluster than the Top1 clusters (**Fig. 4G**). Moreover, when supercoils were immobilized and accumulated, the cluster of Top2A, but not that of Top1, showed an increase proximity to the SMC5/6 cluster (**Fig. 4G**).

These observations are consistent with the idea that SMC5/6 and Top2A are involved in supercoil relaxation in a collaborative manner. To gain further insight into the mechanism by which SMC5/6 and Top2A facilitate supercoil relaxation, we examined the cluster distribution patterns of these molecules at the forks. We found that Top2A clusters tend to localize closer to PCNA than SMC5/6 (**Fig. 4H**), suggesting that Top2A may assemble with supercoils that are confined to a region anterior to that occupied by SMC5/6.

Taken together, our results favor a model in which SMC5/6 confines and stabilizes negative supercoils behind the fork. This, in turn, increases local supercoil density and recruits Top2A to these stabilized supercoils, thereby facilitating their efficient release and subsequent fork progression (**Fig. 5**).

**Figure 5.**
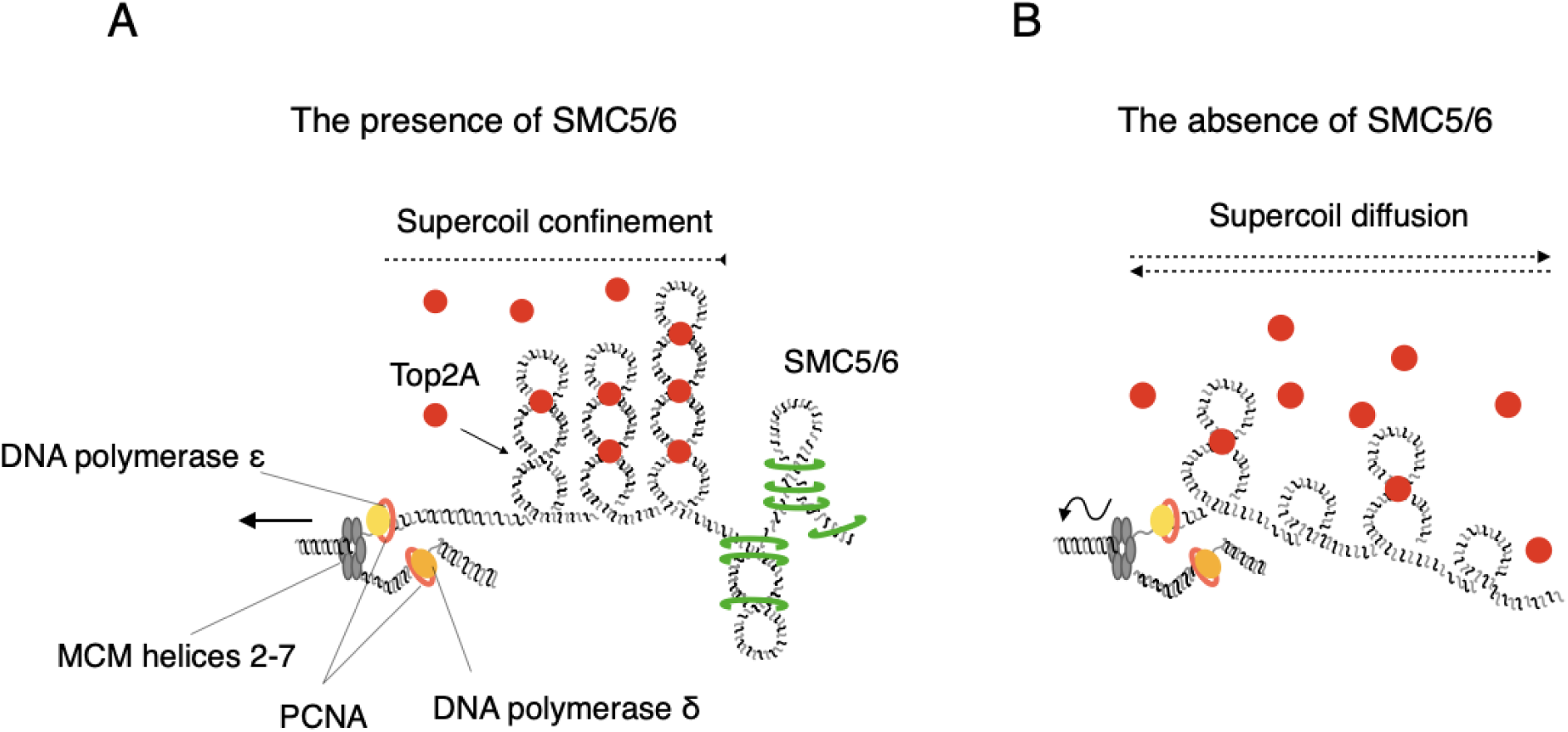
SMC5/6 confines negative supercoils behind the replication fork and promotes Top2 A clustering. Hypothetical model of how SMC5/6 promotes Top2A recruitment to negative supercoils behind the replication fork. Association of SMC5/6 with replicated DNA confines negative supercoils and prevents them from becoming unstable and diffusible. Such confinement increases superhelical density and enhances the density of the Top2A cluster (A). In the absence of proficient SMC5/6, supercoils become unconfined and unstable, reducing the density and increasing the size of negative supercoil and Top2A clusters (B). These differences seem to underlie differences in the cellular ability to handle supercoils during replication.

### Impairment of SMC5/6-mediated enrichment of Top2A in HeLa cells

In light of our understanding of how cells respond to supercoils that are inherently produced during fork progression, and given that replication stress represents a significant source of genomic instability in cancer cells (Burrell *et al*., 2013; Minocherhomji *et al*., 2015; Saxena & Zou, 2022), we postulated that this cellular capacity to resolve supercoils may be susceptible to alterations in transformed cells. In this vein, we found that HeLa cells accumulated more negative supercoils than RPE-1 cells (**Fig. 6A**). The STORM analyses of negative supercoil clusters suggested that HeLa allowed supercoil diffusion to a greater extent than RPE1 cells. This was evidenced by the increased extra-cluster fraction of negative supercoils (**Fig. 6B, 6C**) and the larger negative supercoil clusters with lower density in HeLa cells (**Fig. 6D, 6E**). Furthermore, the STORM analysis of Top2A clusters in HeLa cells showed a molecular distribution comparable to that of negative supercoils (**Fig. 6F-H**). These results suggest that negative supercoils are poorly confined, resulting in insufficient recruitment of Top2A to these supercoils in HeLa cells. We assumed that the enrichment of SMC5/6 is impaired in these cells, which is unlikely to be caused by reduced expression levels of the SMC5/6 (**Fig. EV7A**).

**Figure 6.**
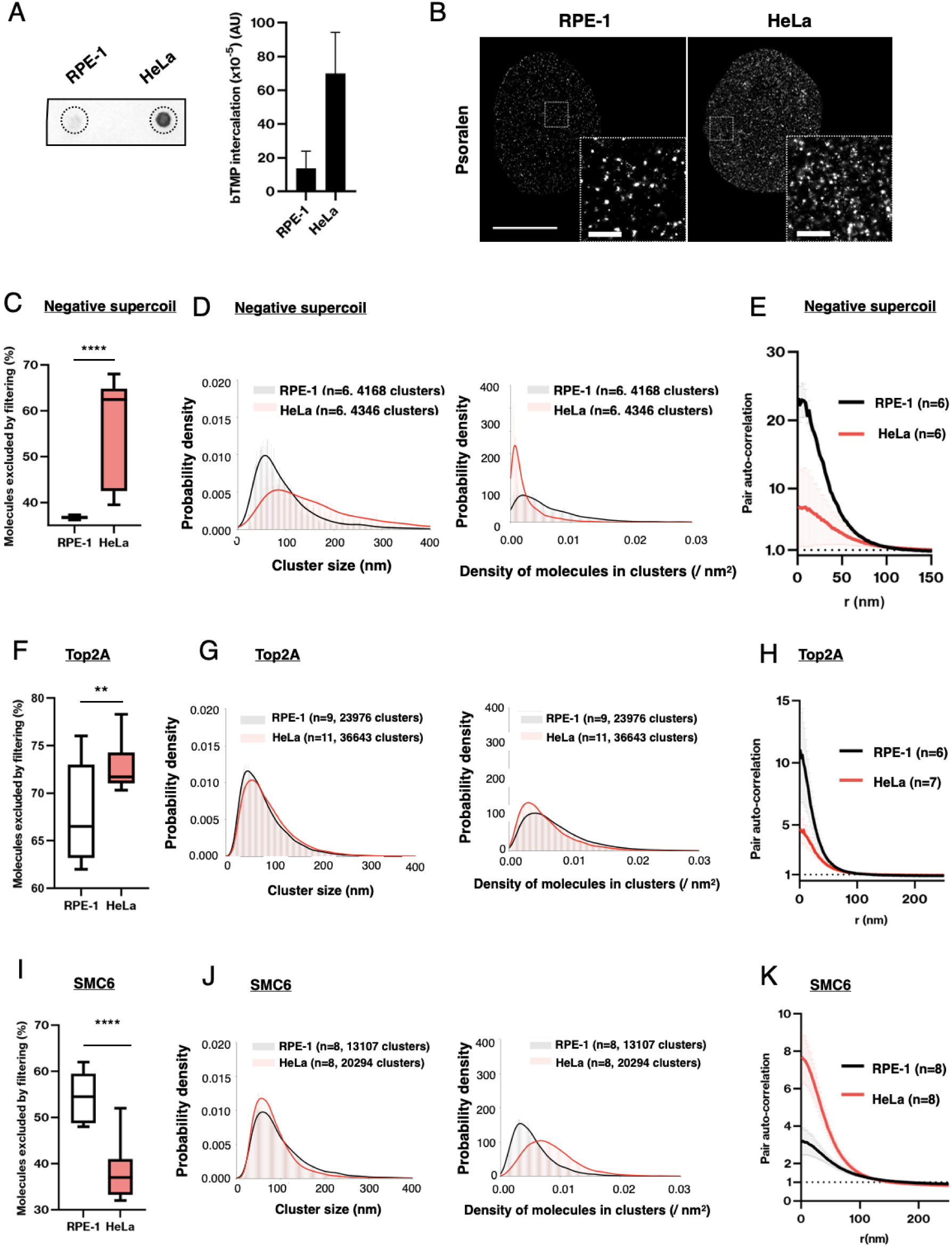
Accumulation and dynamic spatial organization of supercoils in RPE-1 and HeLa cells. (A) Quantitative analysis of negative supercoils. Incorporated bTMP in asynchronous RPE-1 and HeLa cells was intercalated by UV irradiation and then visualized with streptavidin-HRP on extracted and spotted DNAs. Dotted circles indicate where DNA extracts were spotted. Bar charts show signal intensities in arbitrary units (mean ± SD, n=3). (B) Representative STORM images of biotinylated psoralen intercalated in SMC6-degron RPE-1 and HeLa cells. Cells in late S phase were examined by fluorescence microscopy for mCherry-labeled PCNA and by STORM imaging for psoralen molecules, detected with azide-fused Alexa-Fluor-647. Scale bar, 10 μm. Enlarged views show that psoralen molecules are more densely distributed in HeLa cells. Scale bar, 1 μm. (C, D) Segmentation of late S phase nuclei identified ∼6-7 x 10^2^ psoralen clusters per cell in RPE-1 and HeLa cells. Compared with RPE-1 cells, HeLa cells had an increased population excluded from the cluster (C), increased cluster size (D, left) and decreased the molecular density (D, right). Two-tailed Wilcoxon-Mann-Whitney tests indicate differences in the excluded fraction (*P* < 0.0001 shown as ****), the size (*P* < 0.0001), and the density (*P* < 0.0001) between control and IAA. (E) Quantitative analysis of psoralen molecular assemblies in RPE-1 and HeLa cells. (F, G) Quantification of Top2A clusters and molecular assemblies in RPE-1 and HeLa cells. Segmentation of late S phase nuclei identified ∼3 x 10^3^ Top2A clusters per cell. Compared to RPE-1 cells, HeLa cells increased the population excluded from the cluster (F), increased the cluster size (G, left), and decreased the molecular density (G, right). Two-tailed Wilcoxon-Mann-Whitney tests indicate difference in the excluded fraction (*P* < 0.01 shown as **), the size (*P* < 0.0001), and the density (*P* < 0.0001) between RPE-1 and HeLa cells. (H) Pair auto-correlation analysis of SMC6 molecular assemblies in late S phases. RPE-1 cells in black, and HeLa cells in red. Error bars indicate standard deviations. Similar profiles were obtained in three independent experiments. (I, J) Quantification of SMC5/6 clusters and molecular assemblies in RPE-1 and HeLa cells. Segmentation of late S phase nuclei identified ∼3 x 10^3^ SMC6 clusters per cell. Compared to RPE-1 cells, HeLa cells increased the population excluded from the cluster (I), increased the cluster size (J, left), and decreased the molecular density (J, right). Two-tailed Wilcoxon-Mann-Whitney tests indicate difference in the excluded fraction (*P* < 0.0001 shown as ****), the size (*P* < 0.0001), and the density (*P* < 0.0001) between RPE-1 and HeLa cells. (K) Pair auto-correlation analysis of SMC6 molecular assemblies in late S phases. RPE-1 cells in black, and HeLa cells in red. Error bars indicate standard deviations. Similar profiles were obtained in three independent experiments.

Counterintuitively, however, SMC5/6 clusters in HeLa cells had a much higher density and smaller size than in RPE-1 cells (**Fig. 6I-K**). The lower extra-cluster fraction of SMC5/6 in HeLa cells suggested that the majority of SMC5/6 in the nucleus was already allocated to over-accumulated supercoils. To verify this possibility that the dense SMC5/6 clusters in HeLa cells reflect accumulated supercoils, we treated cells with high-dose psoralen and induced additional supercoil accumulation (**Fig. 7A**). This treatment resulted in smaller and denser SMC5/6 clusters in RPE-1 cells (**Fig. 7B, 7C**), and significantly reduced the extra-cluster fraction of SMC5/6 (**Fig. 7D**), supporting the idea that SMC5/6 assembles and increases the density of clusters for accumulated supercoils. Whereas in HeLa cells, SMC5/6 clusters did not grossly accumulate more SMC5/6 molecules (**Fig. 7E**) and the extra-cluster fraction remained unchanged (**Fig. 7F**). These observations suggest that HeLa cells have low redundancy of SMC5/6, making it difficult to supply additional SMC5/6 to newly generated supercoils. This is likely due to the extensive use of nuclear SMC5/6 for over-accumulated supercoils (**Fig. 6I-K**).

**Figure 7.**
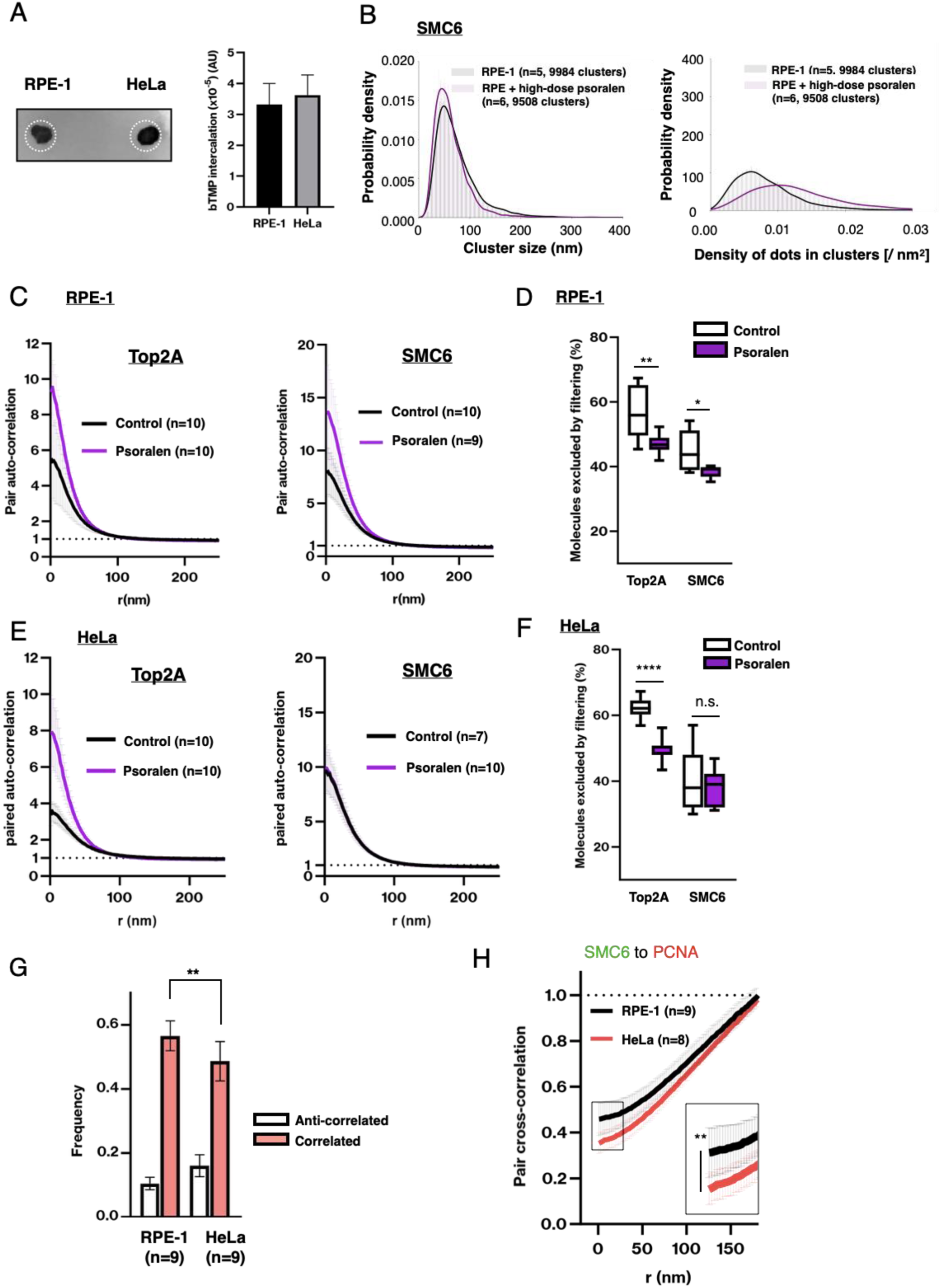
Reduced responsiveness to supercoil accumulation in HeLa cells. (A) Cells were treated with biotinylated psoralen (bTMP) and intercalated bTMP was visualized with streptavidin-HRP on extracted and spotted DNA. White dotted circles indicate where DNA extracts were spotted. Bar charts show signal intensities in arbitrary units (mean ± SD, n=3). (B) Segmentation of late S phase nuclei identified ∼1 x 10^3^ SMC5/6 clusters per cell in RPE-1 cells untreated or treated with high-dose psoralen. Compared to RPE-1 cells, psoralen-treated RPE-1 cells decreased the cluster size (left) and increased the molecular density (right). Two-tailed Wilcoxon-Mann-Whitney tests indicate difference in size (*P* < 0.0001), density (*P* < 0.0001) between RPE-1 and psoralen treated RPE-1 cells. (C) Pair auto-correlation analysis of Top2A and SMC6 molecular assemblies in late S phase of RPE-1 cells. High-dose psoralen treatment increased the density of Top2A assemblies and that of SMC5/6 assemblies in RPE-1 cells. Control in black; high-dose psoralen in purple. Error bars indicate standard deviations. (D) Proportion of excluded fraction of Top2A and SMC6 molecules analyzed in segmentation analysis. RPE-1 markedly reduced the excluded Top2A fraction and the excluded SMC5/6 fraction by high-dose psoralen treatment. **, *P* < 0.01; *, *P* < 0.05; two-tailed Wilcoxon-Mann-Whitney tests. (E) Pair auto-correlation analysis of Top2A and SMC6 molecular assemblies in late S phase of HeLa cells. High-dose psoralen treatment increased the density of Top2A assemblies in HeLa cells, whereas immobilized supercoils did not increase the density of SMC5/6 in HeLa cells. Control in black; high-dose psoralen in purple. Error bars indicate standard deviations. (F) Proportion of excluded fraction of Top2A and SMC6 molecules analyzed in segmentation analysis. RPE-1 and HeLa cells markedly reduced the excluded fraction of Top2A by high-dose psoralen treatment. RPE-1 cells also reduced the excluded SMC5/6 fraction, whereas HeLa cells did not reduce it by psoralen. n.s., not significant; ****, *P* < 0.0001; two-tailed Wilcoxon-Mann-Whitney tests. (G) CBC analysis shows that SMC5/6 is highly correlated with PCNA in late S phase of both RPE-1 and HeLa cells. Anti-correlated distribution, white bars; correlated distribution, red bars. **, *P* < 0.01; two-tailed Wilcoxon-Mann-Whitney test. (H) Correlated distribution of SMC5/6 from PCNA in RPE-1 and HeLa cells. The distribution of SMC5/6 to PCNA is closer in RPE-1 cells than in HeLa cells. **, *P* < 0.01; two-tailed Wilcoxon-Mann-Whitney test was used to compare data points of g(r=0).

By contrast, likewise in RPE-1 cells, supercoils accumulated by psoralen treatment led to an increased density of Top2A clusters even in HeLa cells (**Fig. 7C-F**). Because psoralen intercalates into the supercoils and stabilizes their structures, these results provide further corroborate the idea that stabilization of supercoils by SMC5/6 is required for Top2A clustering on supercoils (**Fig. 5**).

We therefore hypothesized that the insufficient supply of SMC5/6 to the replication forks in HeLa cells causes supercoil diffusion and reduced Top2A recruitment. To validate this, we examined if the spatial distribution of SMC5/6 clusters around replication forks is inferior in HeLa cells compared to RPE-1 cells. The CBC analysis revealed that RPE-1 cells have a higher correlated distribution between PCNA and SMC5/6 (**Fig. 7G**). Moreover, pair cross-correlation analysis indicated that SMC5/6 is localized closer to PCNA in RPE-1 cells than in HeLa cells (**Fig. 7H**). These results are consistent with the interpretation that HeLa cells accumulate many supercoils and therefore require abundant SMC5/6 for their regulation. However, the short of SMC5/6 for newly generated supercoils associated with the forks results in a failure to increase the cluster density of newly generated negative supercoils and Top2A, possibly contributing to supercoil over-accumulation.

## Discussion

The psoralen uptake experiments led us to find that SMC5/6 is required for releasing negative supercoils during S phase. These negative supercoils are thought to arise when DNA polymerases run behind the replication fork (Hingorani & O’Donnell, 2000; Kurth *et al*., 2013; Leu *et al*., 2003; Ullsperger, 1995). Additionally, a fraction of SMC5/6 was found to increasingly associate with chromatin in S phase, which requires the activity of DNA polymerases (**Fig. 1D-F**). Thus, it is plausible that the role of SMC5/6 in releasing supercoils relates to its association with replicated DNA. Rotation of the replication fork can alleviate the accumulation of negative supercoils, but this in turn leads to intertwining of sister chromatids (Kurth *et al*., 2013; Ullsperger, 1995). Taken into account that fork rotation is restricted at fork pausing sites and replication termination sites (Dewar *et al*., 2015; Schalbetter *et al*, 2015; Sundin & Varshavsky, 1980), we consider it likely that a more general function of SMC5/6 in releasing negative supercoils is to stabilize them, at least in human cells.

Quantitative analyses of STROM imaging datasets indicated the presence of clusters of negative supercoils and Top2A in S phase nuclei. Characterizing the geographic distribution of these clusters allowed us to conclude that negative supercoils and Top2A similarly assemble into clusters in an SMC5/6-dependent manner behind the replication fork (**Fig. 2C-H, 3B-F**). Our results showed that SMC6 depletion decreased the density of negative supercoil clusters by increasing cluster size and extracluster fraction (**Fig. 2D-H**). As accumulated negative supercoils behind the progressing fork are locally enriched by SMC5/6, numerous DNA crossovers are generated within these superhelical structures (Boles *et al*., 1990; Vologodskii *et al*., 1992), where Top2 can bind (Zechiedrich & Osheroff, 1990). This results in a concomitant increase in the cluster density of Top2A (**Fig. 5**). In contrast, supercoil diffusion in the absence of SMC5/6 reduces the number of crossovers and decreases Top2A’s cluster density, thereby impeding the efficient Top2A-mediated supercoil relaxation (**Fig. 5**).

How mechanistically does SMC5/6 prevent supercoil diffusion? Single-molecule analysis using magnetic tweezers has shown that increasing the pulling force or torque applied to supercoiled DNA molecules stabilizes supercoil dynamics by preventing the diffusion along the DNA (van Loenhout *et al*., 2012). One possibility is therefore that SMC5/6 exerts a force when it associates with replicated DNA behind the fork (**Fig. 5**). This hypothesis is indeed supported by in vitro experiments, which showed that SMC5/6 is capable of stabilizing supercoils against the force of approximately 1 pN (Gutierrez-Escribano *et al*., 2020; Serrano *et al*., 2020). Although additional torque is shown to impede enzymatic processing of Top2A, a force of ∼1 pN has a negligible effect on its capacity to release supercoils (Strick *et al*., 2000).

Due to the lack of a suitable probe precluded us from analyzing the positive supercoils. However, in vitro studies have demonstrated that SMC5/6 is able to stabilize both positive and negative supercoils (Gutierrez-Escribano *et al*., 2020; Jeppsson *et al*., 2024; Serrano *et al*., 2020), and Top2A is known to relax both supercoils (Ullsperger, 1995; Wang, 2002). It is therefore conceivable that SMC5/6 can also support Top2A-mediated resolution of positive supercoils that arise in front of the fork. Two-color STORM analyses showed that approximately half of the Top2A clusters corresponded to negative supercoils associated with replication (**Fig. 4D**). It seems reasonable to posit that the remaining half may correlate with positive supercoils. SMC5/6 may prevent diffusion of positive supercoils in a manner analogous to its effect on negative supercoils. Additionally, by promoting fork rotation, SMC5/6 can dissipate, and thereby indirectly release, positive supercoils (Dewar *et al*., 2015; Kegel *et al*., 2011; Ullsperger, 1995; Wang, 2002). Together, our observations suggest that SMC5/6 plays a pivotal role in fork progression by releasing supercoiled structures that arise during replication.

HeLa cells accumulated more negative supercoils than RPE1 cells, yet paradoxically revealed reduced cluster densities of negative supercoils and, consequently, of Top2A. It appears as if SMC5/6 is defective in HeLa cells compared with RPE-1 cells (**Fig. 6A-H**), and this is likely due to insufficient supply of SMC5/6 to newly generated negative supercoils (**Fig. 7C-F**). The longer distance observed between the fork and SMC5/6 in HeLa cells (**Fig. 7H**) is consistent with this interpretation. This distance would hamper the ability of SMC5/6 to exert and convey sufficient force behind the fork, or may allow supercoil diffusion backward from the fork, or both (**Fig. 6B-E**). This scenario reasonably explains why HeLa cells cannot form dense clusters of negative supercoils and Top2A, leading to the accumulation of supercoils.

SMC5/6 was found to be highly expressed and assembled as a cellular response to supercoil accumulation in HeLa cells (**Fig. 6I-K, 7B-D**). This observation leads to the question of whether the extensive requirement for SMC5/6 seen in HeLa cells is also present in other cancer cells. Notably, SMC6 expression is markedly elevated in a wide range of human malignancies, with or without recurrences or metastasis, when compared to normal tissues (**Fig. EV7B**). Furthermore, elevated levels of SMC5/6 subunits are observed in 20-60% of cancers across different tumor types (**Fig. EV7C**), and the higher expression levels of SMC6 correlated with a poor prognosis (**Fig. EV7D**). These datasets allow for a previously unappreciated hypothesis: that the spatial organization of supercoil dynamics is disrupted in many cancers. Should this hypothesis be validated, targeting the SMC5/6-Top2A axis could represent a novel therapeutic approach.

## Methods

### Antibodies

The following antibodies were used: SMC6L1 (M01, Cl7one 2E6; Abnova), SMC6 (PA5-30820, Invitrogen), SMC5 (18038; Abcam), NSE4A (AP9909a, abcepta), Top2A (F-12, sc-365916, Santa Cruz), Top2A (ab74715, Abcam), Top1 (A302-589A, Bethyl), CREST sera, α-tubulin (T6074, clone B-5-1-2; Sigma-Aldrich), GFP (ab290; Abcam, IF), histone H2B (ab1790; Abcam), histone H3 (9715S, Cell Signaling Technology), mCherry (GTX 128508; GeneTex), mCherry (M11217, Invitrogen), and GAPDH (14C10, 2118, Cell Signaling Technology). The secondary antibodies used for immunofluorescence (IF) were: Alexa Fluor-conjugated highly cross-absorbed goat anti-rabbit, anti-mouse, or anti-human IgG (H+L) (A-32731, A-11036, A-21245, A-21236, A-11029, Invitrogen). The secondary antibodies used for immunoblots were: horseradish peroxidase (HRP)-labelled Rabbit TrueBlot (18-8816-13, Rockland Immunochemical) and HRP-linked anti-rabbit and mouse IgG (NA9340V, NA9310V, GE Healthcare).

### Cell Culture and lentiviral transfection

HeLa Kyoto cells (gift from Narumiya S., Kyoto University) and RPE-1 cells (ATCC, Cat#CRL-4000) were cultured in DMEM supplemented with 10% fetal calf serum, 0.2 mM L-glutamine, 100 U/ml penicillin, and 100 μg/ml streptomycin at 37 °C in a 5% CO_2_ environment. To generate cells stably expressing cell-cycle reporters, lentivirus encoding DNA helicase B (DHB)-mKO_2_ or -mEGFP, or PCNA-mCherry produced in 293T cells was infected into SMC6-degron HeLa and RPE-1 cells. The construct for DHB expression used in this study is a gift from Sabrina Spencer (Stanford University).

### Generation the cell lines using CRIPSR-Cas9 genome editing

HeLa cells expressing OsTIR1 (wild type) under the Ubc promoter (Ubc-OsTIR1) from AAVS1 locus were generated as described previously (Natsume *et al*, 2016). To generate RPE-1 cells expressing OsTIR1, EF1a-OsTIR1 (F74G) (Yesbolatova *et al*, 2020) coding sequence was cloned into piggyBac plasmid and co-transfected with pCMV-hyPBase vector by electroporation (Neon transfection system, Thermo Fisher Scientific).

SMC6-mAIDmClover, Top1-mCherry, and Top2A-mCherry cell lines tagged at respective endogenous loci were generated by CRISPR-Cas9 mediated homologous recombination using a double-nickase strategy (Koch *et al*, 2018). Briefly, homologous arms surrounding the stop codons of SMC6, Top1, and Top2A were cloned into pGEMTeasy vector (Promega). mAIDmClover-Neomycin, mAIDmClover-Hygromycin, mAIDmClover-Blasticidin, and mCherry-Blasticidin coding sequences were introduced before the stop codon. These homology-directed recombination donor plasmids were transfected to cells with pairs of CRISPR short guide RNAs (sgRNAs), which are targeted at their C termini. These plasmids were transfected to HeLa cells using Fugene HD (Promega) and to RPE-1 cells using electroporation (Thermo Fisher Scientific). mAIDmClover or mCherry-positive single cells were isolated by FACS soring (Aria Ⅲ, Becton Dickinson) after two-week treatment of hygromycin, G418, or blasticidin. Homozygous targeting of genomic alleles was confirmed by genomic PCR and Western blotting. To deplete SMC6, HeLa cells and RPE-1 cells were cultured in the presence of 500 μM IAA or 1 μM 5Ph-IAA, respectively. The constructs for CRISPR-Cas9 mediated gene editing and piggyBac system and the synthetic IAA/5Ph-IAA were gifts from Masato Kanemaki (National Institute of Genetics, Japan).

### Cell synchronization, inhibitor treatment, and S-phase release experiments

To synchronize SMC6-degron HeLa cells at G1/S boundary, cells were treated with 2 mM thymidine (T1895–1G; Sigma-Aldrich) for 16 h, and then released after washing once with DMEM. Nine hours later, cells were treated with 1 μg/ml aphidicolin (011-09811, Wako) plus 33 μM XL-413 (BMS-863233, AdooQ BioScience). To degrade SMC6 prior to G1/S release, IAA was supplemented with aphidicolin plus XL-413. For S-phase release experiments, SMC6-degron HeLa cells arrested at G1/S were released into S phase for the indicated period of time (0-9 h) and then, the cells were treated with 10 μM EdU for 20 min. To analyze replication under Top1 or Top2 inhibition, 10 nM camptothecin (Wako) or 100 nM ICRF-193 (Sigma-Aldrich) were added upon G1/S release. To assess the progression of S phase of SMC6-degron RPE-1 cells, cells synchronized with the 150 nM Cdk4/6 inhibitor palbociclib (Selleck) for 24 h in G1 were released into the first S phase with or without 5Ph-IAA.

### Immunofluorescence and click chemistry microscopy

Cells were fixed with 4% paraformaldehyde (PFA) in 0.137 M sodium phosphate buffer (pH 7.4) for 15 min. Fixed cells were permeabilized with 0.2% Triton-X in PBS for 10 min and incubated with 3% bovine serum albumin (BSA) in 0.01% Triton-X in PBS (PBS-T) for 30 min. Cells were incubated with primary antibodies for 1 h at room temperature or overnight at 4°C, and incubated with secondary antibodies for 40 min at room temperature. After counterstaining DNA by 5 min incubation of 0.1 μg/ml DAPI, coverslips were mounted with Prolong Gold anti-fade mounting reagent (Invitrogen). Incorporated EdU was detected with azide-fused Alexa-Fluor-647 or azide-fused Alexa-Fluor-488 (for STORM) by the Click-iT^TM^ Plus Kit (C10640, C10637; Thermo Fisher). Images were acquired with a Zeiss wide field fluorescence microscope, 63x objective lens and a PRIME BSI camera (Photometrics), or a confocal LSM880 microscope (Zeiss) and 63x objective lens with Airyscan detection. For high-content quantitative analysis of EdU fluorescence intensities, fluorescence images were acquired with 10x objective lens of the high-content imaging systems (Operetta CLS, PerkinElemer or CQ1, Yokogawa) and processed for the semi-automated analyses (Harmony, PerkinElemer or CellPathfinder, Yokogawa). Briefly, nuclei were identified as round DAPI-positive areas and the mean EdU intensity per nucleus was measured. For the classification of early, middle, and late S phases, the difference in incorporated EdU appearances between three S phases was distinguished by machine learning based analysis using the CellPathfinder software (Yokogawa).

### Chromatin fractionation

Briefly, to obtain chromatin-enriched fraction, cells were resuspended (4 × 10^7^ cells/ml) in buffer A (10 mM HEPES, pH 7.9, 10 mM KCl, 1.5 mM MgCl_2_, 0.34 M sucrose, 10% glycerol, 1 mM dithiothreitol [DTT], a protease inhibitor cocktail [Roche]). Triton X-100 (0.1%) was added, and the cells were incubated for 5 min on ice. Nuclei were pelleted by low-speed centrifugation (4,000 rpm for 5 min in 4°C). The supernatant containing the cytoplasmic fraction was further clarified by high-speed centrifugation (15,000 rpm for 30 min in 4°C) to remove cell debris and insoluble aggregates. The precipitate was washed once and then resuspended in buffer A containing Triton X-100 (0.1%) and verified by immunoblotting (Mendez & Stillman, 2000).

### Immunoblot and immunoprecipitation

Cell pellets were lysed in immunoprecipitation buffer (20 mM Tris-HCl, pH 7.5, 150 mM NaCl, 20 mM β-glycerophosphate, 5 mM MgCl_2_, 0.1% NP-40, 1 mM DTT) containing a protease inhibitor cocktail and 0.05% Omnicleave^TM^ endonuclease (OC7850K, cambio) on ice for 20 min. Cell extracts, after the insoluble fraction was removed by centrifugation at 15,000 rpm for 30 min at 4°C, were used for immunoprecipitation. Total protein concentrations of cell lysates were measured by Bradford-basis method (Protein Assay System, Bio-Rad Laboratories) and the loading protein amounts per lane were controlled. The cell extracts were resolved by SDS–PAGE and transferred to a PVDF membrane (Immobilon-P, Millipore). Can Get Signal Immunoreaction Enhancer Solution 1 (Toyobo) was used to dilute antibodies. For immunoprecipitation, supernatant was incubated with Surebead Protein G Magnetic beads (Bio-Rad) coupled to anti-GFP antibodies at 4°C for 30 min, washed three times with immunoprecipitation buffer, and then suspended in SDS sample buffer.

### Chromosome spreads

Mitotic cells were collected by shaking off, 30-45 min after the release from the G2 arrest after 9 μM RO-3306 treatment for 3-4 h (ALX-270-463-M005, Enzo Life Science) or 2.5 h synchronization by 100 ng/ml colcemid (AG-CR1-3567-M005, Funakoshi). To assess MiDAS, G2 arrested cells were released into medium containing 10 μM EdU. To inhibit transcription, cells were treated with 1 μM triptolide (T2899, TCI Chemicals) during G2 arrest and subsequent release into mitosis. After 5 min incubation with a hypotonic buffer (PBS : H_2_O = 1:4), cells were fixed with fresh Carnoy’s solution (70% methanol, 30% acetic acid). Fixed cells were dropped on glass slides and dried. For staining chromosomes, slides were stained with 5% Giemsa solution for 5 min, washed with water for 3 times, air-dried, and mounted with Entellan embedding agent (107961, Sigma-Aldrich).

### Fluorescence in situ hybridization

Labelling telomeres and centromeres were conducted as previously described (Lansdorp *et al*, 1996). Chromosomes spread on glass slides were fixed with 4% formaldehyde in PBS for 2 min, treated with 1 mg/ml pepsin in 37 mM HCl for 15 min at room temperature, washed with PBS briefly and were dehydrated with a series of 5-min 70%, 90% and 100% ethanol washes and air dried. Hybridization mixture containing Cy3-conjugated (CCCTAA)_3_ oligonucleotide probe and FAM-conjugated a-satellite DNA probe was added to the sample and mounted with coverslip. DNA was denatured by heat at 85°C for 5 min. After hybridization for 2 h at room temperature in the dark, DNA was stained with DAPI at 0.1 μg/ml for 5 min.

### Spectral karyotyping analysis

Probes for spectral karyotyping were purchased from MetaSystems (D-0125-120-DI, 24XCyte) and spread chromosomes were stained following manufacturer’s instruction. Image scan and acquisition were performed with a Axio Imager Z2 microscope (Zeiss) and Plan-Apochromat 10x or 63x lens. Acquired images were analyzed with an Isis imaging software (MetaSystems). To analyze the relationship between the chromosome length and the number of chromatid gaps, chromosome length information for somatic and X chromosomes was obtained from the UCSC Genome Browser (https://genome.ucsc.edu/index.html).

### Fluorescence-activated cell sorting analysis

Cells were harvested by trypsinization, fixed with 70% ethanol precooled at −30°C and incubated at least for 30 min at −30°C. DNA was stained with 50 μg/ml propidium iodide solution and 100 μg/ml RNase A and subjected to flow cytometric analysis on FACSCalibur using Cell Quest (Becton Dickinson) software.

### Fluorescent Recovery After Photobleaching

Living cells expressing both DHB-mKO2 and mClover-SMC6 or both DHB-mEGFP and Top2A-mCherry cultured in CO_2_-independent medium without phenol red (Life Technologies-BRL) were subjected to FRAP analysis as previously described (Christensen *et al*., 2002; Gerlich *et al*, 2006), using Zeiss LSM 880 inverted confocal laser scanning microscope equipped with a CO_2_-controlled on-stage heating chamber and heated 40x water-immersion objective lens. Serial images before and after the bleaching with 0.135 s time intervals were acquired. An nuclear area of 1.5 x 15 μm in size were photobleached with three iterations using 100% of 488 nm and 10% of 405 nm laser powers for SMC6-mClover and 100% of 568 nm and 10% of 405 nm for Top2A-mCherry. Fluorescence intensities of the bleached and unbleached regions were measured for each time point and corrected for extracellular background intensity and for the overall loss in total intensity, caused by the bleach pulse and multiple scans.

### STORM imaging analysis

The immunostained SMC6-degron HeLa and RPE-1 cells on the cover glass were mounted with freshly prepared oxygen scavenger imaging buffer (50 mM Tris-HCl pH 8.0, 10 mM NaCl, 10% glucose, supplemented with 0.7 μg/μL glucose oxidase (Sigma, #G2133-250KU), 0.04 μg/μL catalase (Fujifilm, #035-12903) and 70 mM 2-mercaptoethanol) and set on the glucose oxidase enzymatic system (GLOX) for STORM using a frame-sealed incubation chamber (Bio-Rad). 2D-STORM image acquisitions were performed using a Nikon N-STORM with Eclipse Ti-E inverted microscope and laser TIRF illuminator (Nikon Instruments Inc.). For single-color STORM, Alexa 647 fluorophores were stochastically excited using the 640 nm laser beam with an additional weak pulse of 405 nm laser. For double-color STORM, excitation of 640 fluorophores was followed by that of Alexa 541 fluorophores using the 541 nm laser beam, with an additional 405 nm weak pulse. Images were acquired with an Andor iXon 897 EMCCD camera (Andor Technologies) and a CFI Apo TIRF 100× objective lens (N.A. 1.49). A stack of ∼10,000 frames was acquired with an exposure time of 16 ms for each color of each cell. A compile of the two dimensional coordinates of the detected blinking spots was extracted and corrected for drifts using Nikon NIS-Element imaging software (Nikon Instruments Inc.). To characterize molecular distributions, we applied two methods: the Hierarchical Density-Based Spatial Clustering of Applications with Noise (HDBSCAN) clustering and cropped a region of the cell nucleus as the convex hull of clustered dots, and the pair-correlation statistics of the compiled coordinates (*x_n_*, *y_n_*), in which the density of the molecular fluorescent bright dots becomes *ρ* = *N*/*A* [μm^−2^], where *N* is the number of the coordinates, and *A* [μm^2^] is the area of the nucleus region, as detailed below.

### Clustering analysis by Voronoi filtering and HDBSCAN

To compare 2D-STORM data of different cells in the same condition, we first fixed the density ρ in the cell nucleus and randomly sampled the coordinates of molecular fluorescent bright dots for each cell. Then, we filtered the coordinates according to the local density based on the Voronoi diagram. The Voronoi segmentation algorithm in SciPy (Virtanen *et al*, 2020) allows for calculating the localization density for each coordinate of the dots. When we also calculate the average local density, *δ*, for uniformly distributed dots in the same nucleus region, the value of the twice becomes a good measure as a threshold of spatially uniform random distribution (Levet *et al*, 2015). Then, we normalized the localization density of each dot by *δ* and filtered the dots less than 2*δ*. We can plot the filtered dots with a color map. We also count the rate of how many dots are excluded by the filtering. In addition, since the filtered dots are located forming clusters, we used Hierarchical DBSCAN (HDBSCAN) (Pedregosa *et al*, 2011) to find clusters. The HDBCSAN algorithm requires only one parameter, the minimum number of samples in a group for that group to be considered a cluster, and we set it to 10. Then, we enclosed every detected cluster by the convex hull of dots in a cluster and calculated the cluster size as the root square of the area of a cluster and the density of molecules in clusters.

### Coordinate-based colocalization analysis

To estimate the degree of colocalization for the 2-color 2D-STORM data, we calculated the coordinate-based colocalization (CBC) values (Malkusch *et al*., 2012; Pageon *et al*., 2016). Let {*A_i_*} and {*B_i_*} represent the lists of coordinates from two channels. The CBC value *C_A_i__* is assigned to each coordinate *A_i_*as a value ranging from −1 to 1. The central part of the calculation involves comparing the distributions of {*A_i_*} and {*B_i_*} around a coordinate *A_i_*within a maximum radius *R*_max_ using Spearman’s correlation coefficient. We implemented the original definition by Malkusch *et al*. into our Python code, fixing the density of coordinates in both channels by randomly selecting the same number of coordinates, and setting *R*_max_ to 400 nm with a binning width of 10 nm.

### Pair auto-correlation analysis of dots in the cell nucleus

The pair auto-correlation function is one of the two-point distance statistics based on information from the distance matrix *r_ij_*. When we deal with *O*(10^5^) coordinates, the data size of the two-point distances becomes *O*(10^10^) and is inadequate for a realistic calculation. Therefore, we introduced an upper-distance *R* and focused on the coordinates within the circle region around the *n*th coordinate. Then, we can define the pair auto-correlation function around the *n*th coordinates as

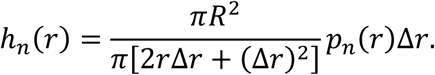

Here, *p_n_*(*r*)Δ*r* corresponds to the existence probability of the dots within the ring region between the radius from *r* to *r*+Δ*r* around the *n*th dots. By averaging for all coordinates, we can calculate the pair auto-correlation function for *O*(10^5^) molecular localizations:

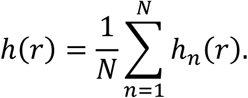

Furthermore, we eliminated the boundary effect of *h*(*r*). Generally, the pair auto-correlation function for the uniformly random distributed dots *h*_random_(*r*) becomes the unit value for any *r*. However, in the finite cell nucleus region, *h_n_*(*r*) of the dots near the boundary inevitably includes a bias due to the location. Therefore, we generated the same number of random dots obeying the uniform distribution in the nucleus region as in calculating *h*(*r*). Then, we computed *h*_random_(*r*) for the coordinates and finally obtained the pair auto-correlation function by this equation:

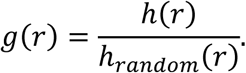

Note that the value of *g*(*r*) depends on the upper-distance *R* and the density of the dots *ρ*. Here, we fixed *R* = 400 nm. For comparing *g*(*r*) in various cell conditions, we also tuned the dot density by randomly selecting the coordinates from the data list. The value of fixed density *ρ*_fixed_ is specified in each analysis. The *ρ*_fixed_ values used include the following: 190 (Fig. 2H), 1000 (Fig. 3F, 3G), 500 (Extended Data Fig. 4G), 700 (Extended Data Fig. 4H), 500 (Fig. 4G), 200 (Fig. 4H), 150 (Fig. 6E), 300 (Fig. 6H), 800 (Fig. 6K), 400 (Fig. 7C, 7E, SMC6), 500 (Fig. 7C, 7E, Top2A), and 400 (Fig. 7H). Besides, we used the binning width Δ*r* as 2 nm.

### Pair cross-correlation analysis of dots in the cell nucleus

When we conduct the 2-color 2D-STORM imaging, the output data includes the list of coordinates from two channels. Then, we can consider four types of pair correlation functions: *g*_ch1 to ch1_(*r*), *g*_ch2 to ch2_(*r*), *g*_ch1 to ch2_(*r*) and *g*_ch2 to ch1_(*r*). The two formers correspond to the auto-correlation of the two channels, and the two latter are the cross-correlation from the first channel to the second channel and vice versa. When we focus on the spatial distribution of the coordinates of the first channel to the second channel, we have to calculate *g*_ch1 to ch2_(*r*) and *g*_ch2 to ch2_(*r*). Then, the ratio *g*_ch1 to ch2_(*r*)/*g*_ch2 to ch2_(*r*) becomes a normalized measure of how the molecules detected from the first channel are relatively distributed to the molecules of the second channel. We fixed the density of coordinates in both channels and randomly selected the same number of coordinates to calculate the pair cross-correlation *g*_ch1 to ch2_(*r*). Calculating the existence probability of dots of the molecule in the first channel for the *n*th coordinate of the second channel, we finally obtained the values of *g*_ch1 to ch2_(*r*). Here, we also fixed the upper-distance *R* = 400 nm and the binning width Δ*r* = 2 nm.

### Live-cell imaging analysis

Cells were suspended in DMEM in 24 well glass bottom plate (5826-024, Iwaki). DNA was stained with siR-DNA (CY-SC007, Cytoskeleton). Images were captured every 1 min, with 100 ms exposure times, through a 40x objective lens (Operetta CLS, PerkinElmer) or 60x objective lens (CellVoyger CQ1, Yokogawa) in high-content image systems. A series of projected images of seven Z-sections with 9 μm intervals were analyzed. For data analysis, images were processed using Fiji (http://fiji.sc.).

### Detection and immobilization of DNA supercoils

Fluorescence detection of incorporated bTMP and immobilization of supercoils in in STORM analysis were performed as previously described (Naughton *et al*., 2013). Briefly, cells grown on coverslips were treated with 1 mg/ml bTMP (Thermo Fisher) in PBS for 20 min at room temperature in the dark. For cluster analysis of negative supercoils, cells were incubated with 10-20 μg/ml bTMP on ice for 2 min in the dark. bTMP was then cross-linked to the DNA by 360 nm UV irradiation for 2∼10 min. Cells were fixed with 4% PFA for 15 min followed by permeabilization with 0.2% PBS-T for 10 min. For staining of biotinylated psoralen, samples were incubated with Alexa fluor 568- or 647-conjugated streptavidin (S11226, S21374, Thermo Fisher) for 1 h at room temperature.

Changes in the amount of bTMP incorporated into cellular DNA were detected as previously described (Aze *et al*, 2016; Naughton *et al*., 2013) with modifications. Samples were incubated with 10 μg/ml bTMP on ice for 5 min in the dark and irradiated with 365 nm UV light for 5 min on a precooled metal block or left untreated. The procedure from addition of psoralen to irradiation with UV light was repeated three times. In other experiment, cells were treated with high-dose psoralen, 1mg/ml bTMP in PBS for 30 min at room temperature in the dark and irradiated for 10 min (**Fig. 5A**). Genomic DNA was extracted from the cells using QIAamp DNA Blood Mini Kit (51104, QIAGEN). RNase A (30141-14, Nacalai tesque) was added to samples with protease during the extraction process. After being fragmented by sonication (Biruptor Ⅱ), the concentration of the extracted DNA samples was measured using Qubit dsDNA HS assay kit (Q32851, Thermo Fisher) and Qubit 4 fluorometer (Q33238, Thermo Fisher). Biotin incorporation into DNA was detected by dot blotting using streptavidin-HRP (Invitrogen) as a probe.

## Acknowledgement

We thank Masato Kanemaki (National Institute of Genetics, Japan) and Sabrina Spencer (Stanford University, USA) for reagents, Utako Kato (JFCR, Japan) for technical assistance, Yasukazu Daigaku (JFCR, Japan) and all members of the T.H. laboratory for discussions. Research in the T.H. lab is supported by the Japan Society for the Promotion of Science (JSPS) Grant-in-Aid for Scientific Research (22H04996, 22H00458, 18H04034 [to T.H.]). Y.K. acknowledges the support from the JSPS Fellowship (DC2).

## Author contributions

Y.K. and T.H. conceived the study. Y.K. designed and performed all cell biological experiments and all STORM image acquisition. S.S. and S.O. performed formal analysis of STORM images. R.N. contributed to some stable cell establishment. K.S. and S.T. performed SKY analysis. Y.K. and T.H. wrote the paper with input from all other authors. T.H. supervised the project.

---

**Figure EV1.**
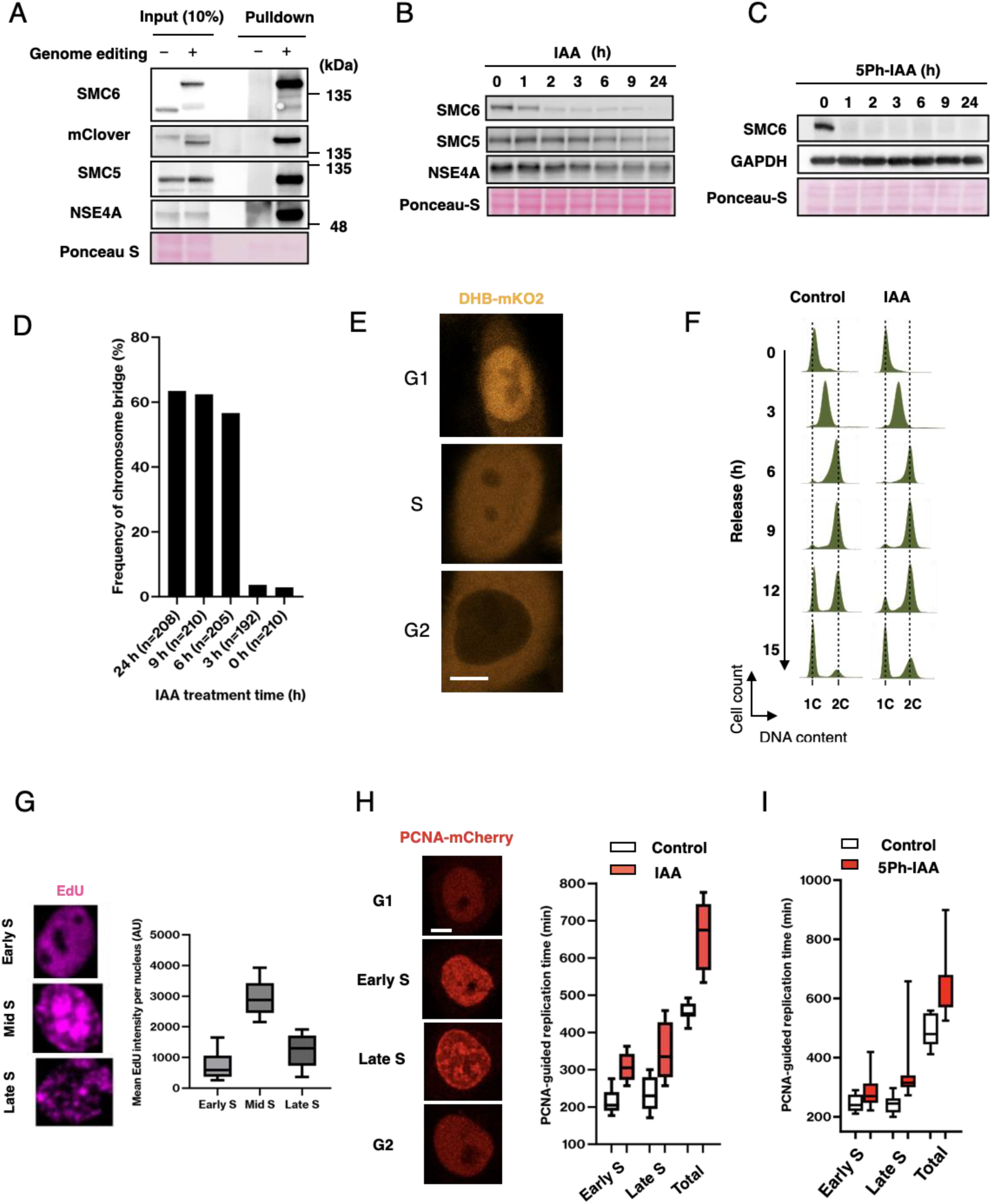
SMC5/6 is required to complete DNA replication prior to mitosis. (A) Tagging of SMC6 with mAID-mClover does not disrupt complex formation. SMC6 endogenously tagged with mAID-mClover was expressed in HeLa cells (SMC6-degron HeLa cells) and immunoprecipitated with anti-GFP antibody, which precipitated mClover-tagged SMC6 with the core subunits of the SMC5/6 complex, SMC5 and NSE4A. (B) Rapid degradation of SMC6 in SMC6-degron HeLa cells after IAA addition. Note that SMC5 and NSE4A were also become unstable by the degradation of SMC6. (C) Rapid degradation of SMC6 in SMC6-degron RPE-1 cells. Total cell extracts were examined at the indicated time points after 5Ph-IAA addition. Note that the faster degradation kinetics of SMC6 in RPE-1 degron cells did not exhibit anaphase bridges at earlier timepoints than observed in HeLa cells (Figure 1C). (D) Bar charts show the frequency of anaphase bridges after SMC6 depletion in HeLa cells. SMC6 was degraded in: G2 (for 3 h treatment), late S (6 h), early S (9 h) and G1 (24 h) phases. (E) In FRAP analysis, stably expressed DHB-mKO_2_ was used as a cell cycle reporter, which allows to distinguish G1, S, and G2 phases, as indicated (Spencer *et al*, 2013). DHB-mKO_2_, orange; SMC6-mClover, green. (A) Flow cytometric analysis of SMC6-degron HeLa cells. The first round of replication progression from the G1/S boundary after the SMC6 depletion is shown. (G) Representative images of incorporated EdU for early, mid, and late S phases, which we used as training datasets for machine learning cell categorization (left). Mean EdU fluorescence intensities for ∼1,500 cells were measured for each S phase and shown in box plots (right). Note that each S phase stage can be differentiated by the incorporated pattern and is distinguishable by the fluorescence intensities of incorporated EdU. (H) Time-lapse imaging of HeLa cells expressing mCherry-PCNA. Representative images of mCherry-PCNA for indicted phases (left). The lengths of replication time in each replication domain are shown based on tracking the mCherry-PCNA (right). IAA was added 2 h prior to tracking. Control, white; IAA, red. Scale bar, 10 μm. (I) Time-lapse imaging of RPE-1 cells expressing mCherry-PCNA. The measured duration of early, late, and total S phase is shown in box plots. 5Ph-IAA was added 2 h before the imaging. Control, white; 5Ph-IAA, red. Scale bar, 10 μm.

**Figure EV2.**
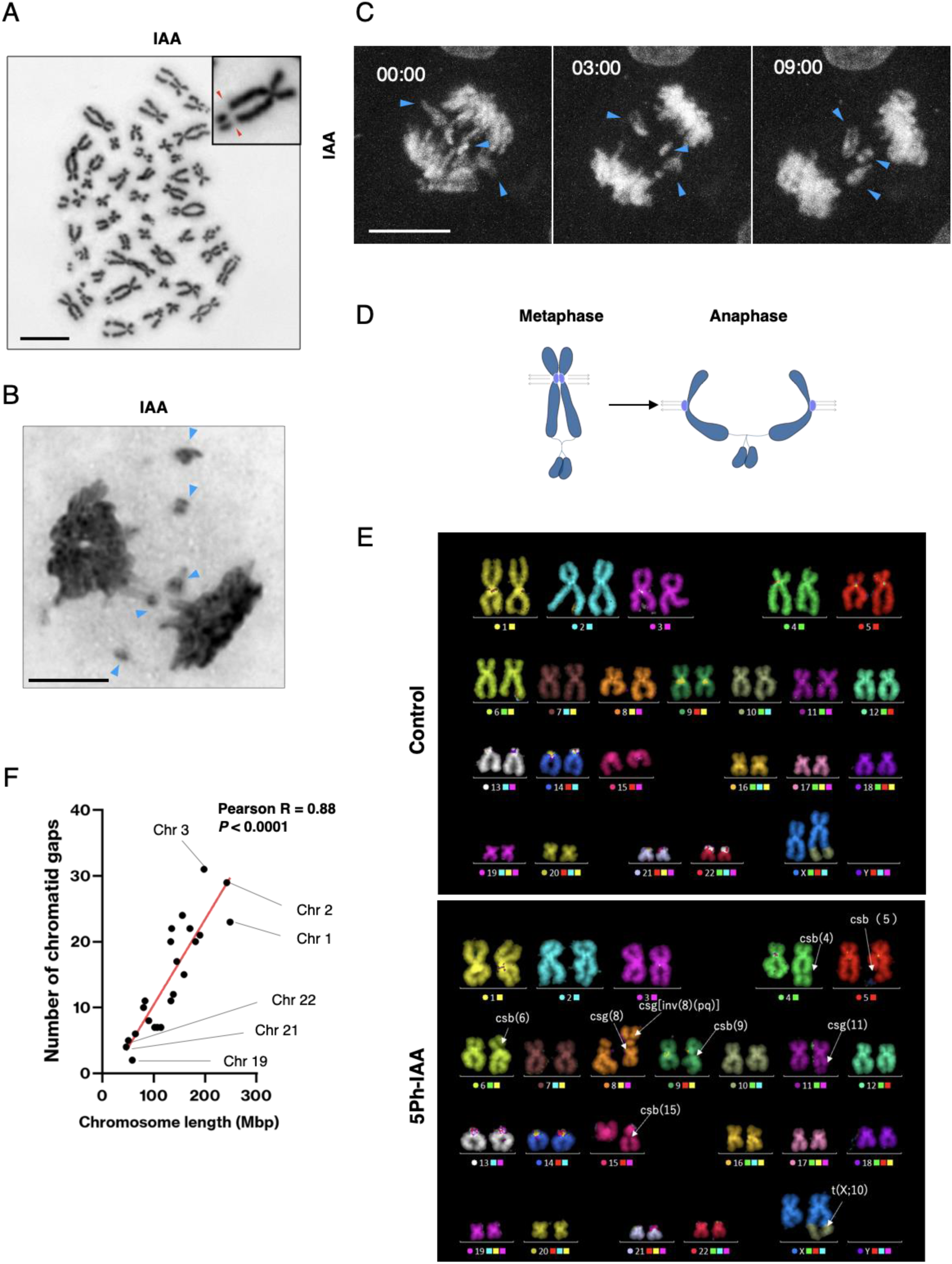
Incomplete DNA replication in the absence of SMC5/6 leads to anaphase bridge. (A) Giemsa-stained spread metaphase chromosomes from SMC6 depleted cells. A magnified view shows a representative chromosome with symmetric chromatid gaps (red arrowheads). Scale bar, 10 μm. (B) Giemsa-stained anaphase chromosomes from SMC6 depleted cells. Small chromatid fragments lagging between the bulk of sister chromatids (blue arrowheads) were frequently observed. Scale bar, 10 μm. (C) Time-lapse imaging analysis showing that the paired chromatid fragments (blue arrowheads) on the chromosome bridges remained between the separating sister chromatids in anaphase. DNA, gray. Scale bar, 10 μm. (D) Symmetric chromatid gaps in metaphase give rise to paired chromatid fragments on the anaphase bridge in cells depleted of SMC6. (E) Spectral karyotyping analysis of the SMC6-degron RPE-1 cells before (Control) or after (5Ph-IAA) SMC6 degradation. Note that in the lower, SMC6-depleted sample, multiple abnormalities in chromosome structure are found, including chromatid gaps (csg) and chromosome breaks (csb) are found. The reported translocation between chromosome X and 10, t(X;10), in RPE-1 cells (Chadwick & Willard, 2003) is also detected. Correlation of the number of chromatid gaps with chromosome length. Each dot represents different chromosomes. Pearson’s correlation coefficient R = 0.88.

**Figure EV3.**
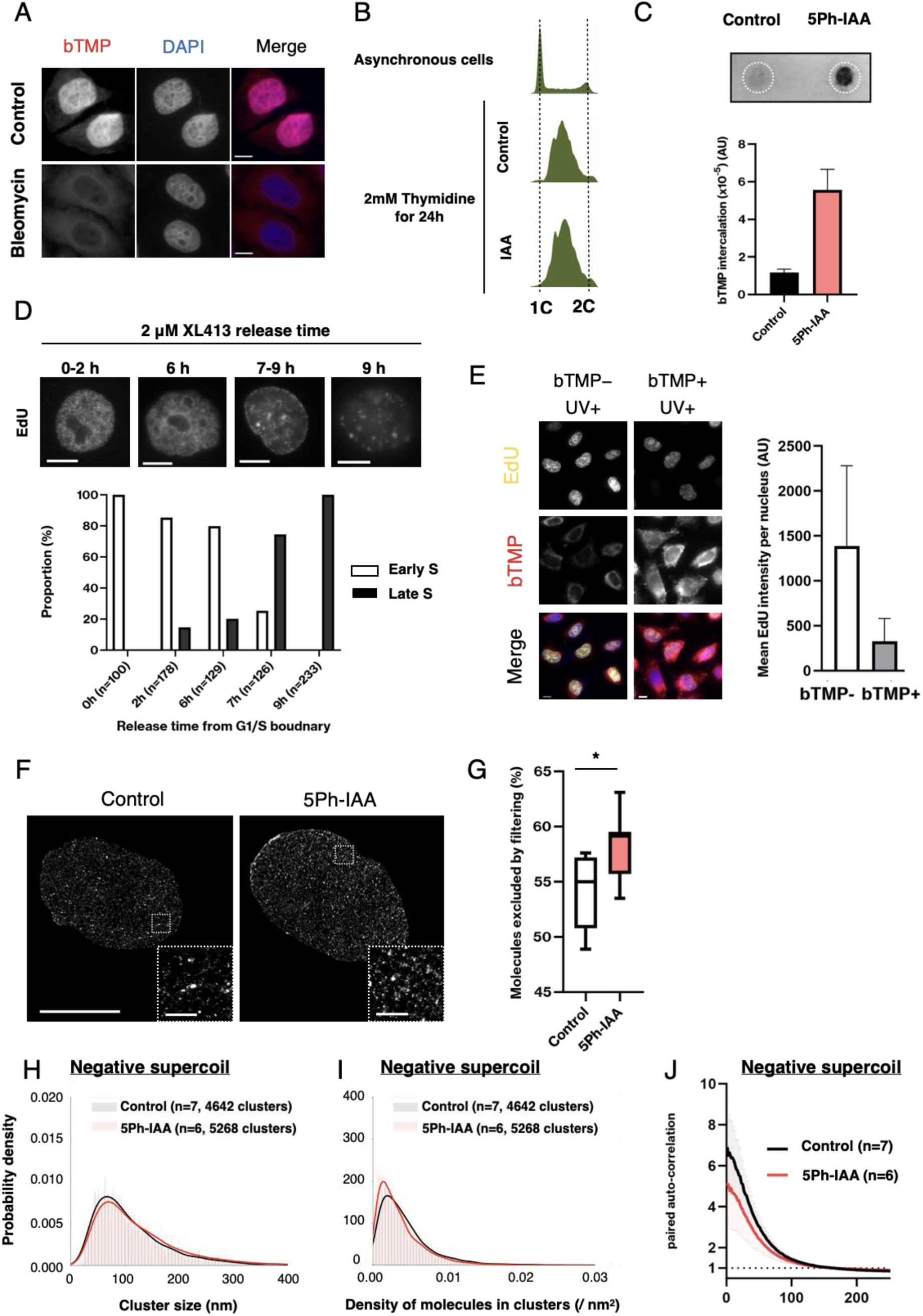
Unconstrained negative supercoils in the absence of SMC5/6 perturbs replication. (A) SMC6-degron HeLa cells with or without 30 min of bleomycin treatment were treated with biotinylated psoralen (bTMP) and intercalated bTMP was visualized with Alexa-conjugated streptavidin. Note that bleomycin treatment eliminated supercoils from nuclei beyond the detection level (bottom panel). Scale bar, 10μm. (B) Flow cytometric analysis of DNA content for asynchronous or late S phase-synchronized SMC6-degron HeLa cells in the presence or absence of IAA. Cells were treated with 2 mM thymidine for 24 h to enrich them in S phase. (C) SMC6-degron RPE-1 cells synchronized in S phase with or without 5Ph-IAA were treated with bTMP and intercalated bTMP was visualized with streptavidin-HRP on extracted and spotted DNA. White dotted circles indicate where DNA extracts were spotted. Bar charts show signal intensities in arbitrary units (mean ± SD, n=3). (D) Enrichment of early S and late S phase populations using a low-dose CDC7 inhibitor XL-413, by inhibiting origin firing of late replication domains (Upper). SMC6-degron HeLa cells arrested at G1/S boundary were released in the presence of 2 μM XL-413 for the indicated period of time and then briefly treated with EdU to differentiate early and late S phases. Representative images of EdU-stained cells at each release timepoint. Cells in early S phase dominate until 6 h after the release, whereas cells in late S phase dominate from 7 h on (Lower). Scale bar, 10 μm. (E) Immobilization of supercoils inhibits DNA replication. SMC6-degron HeLa cells with or without bTMP intercalation were briefly treated with EdU. EdU fluorescence intensities were analyzed for at least 1,000 cells each. Representative immunofluorescence images that were measured for the nuclear EdU intensities. Scale bars, 10 μm. (F) Representative STORM images of biotinylated psoralen incorporation in SMC6-degron RPE-1 cells. Cells in late S phase were analyzed by fluorescence microscopy for mCherry-labeled PCNA and by STORM imaging for psoralen molecules, detected with azide-fused Alexa-Fluor-647. Scale bar, 10 μm. Enlarged views show that psoralen molecules are densely distributed in the SMC6-depleted cell nucleus. Scale bar, 1 μm. (G-I) Segmentation of late S phase nuclei identified ∼7-8×10^2^ psoralen clusters per cell and in control and 5Ph-IAA treated RPE-1 cells. The proportions of psoralen molecules excluded from the clusters are shown in box plots (G). * indicates *P* < 0.05; two-tailed Wilcoxon-Mann-Whitney test. Diagrams show the probability density of the cluster size (H) and the cluster density (I). The fitted mean curves for control and 5Ph-IAA treated cells are shown in black and red, respectively. Two-tailed Wilcoxon-Mann-Whitney tests indicate differences in the size (*P* < 0.0001) and density (*P* < 0.0001) between control and 5Ph-IAA. (J) Quantitative analysis of psoralen molecular assemblies in RPE-1 cells. Densities of psoralen assemblies in late S phase are analyzed in SMC6 depleted (5Ph-IAA, red) and control (black) cells.

**Figure EV4.**
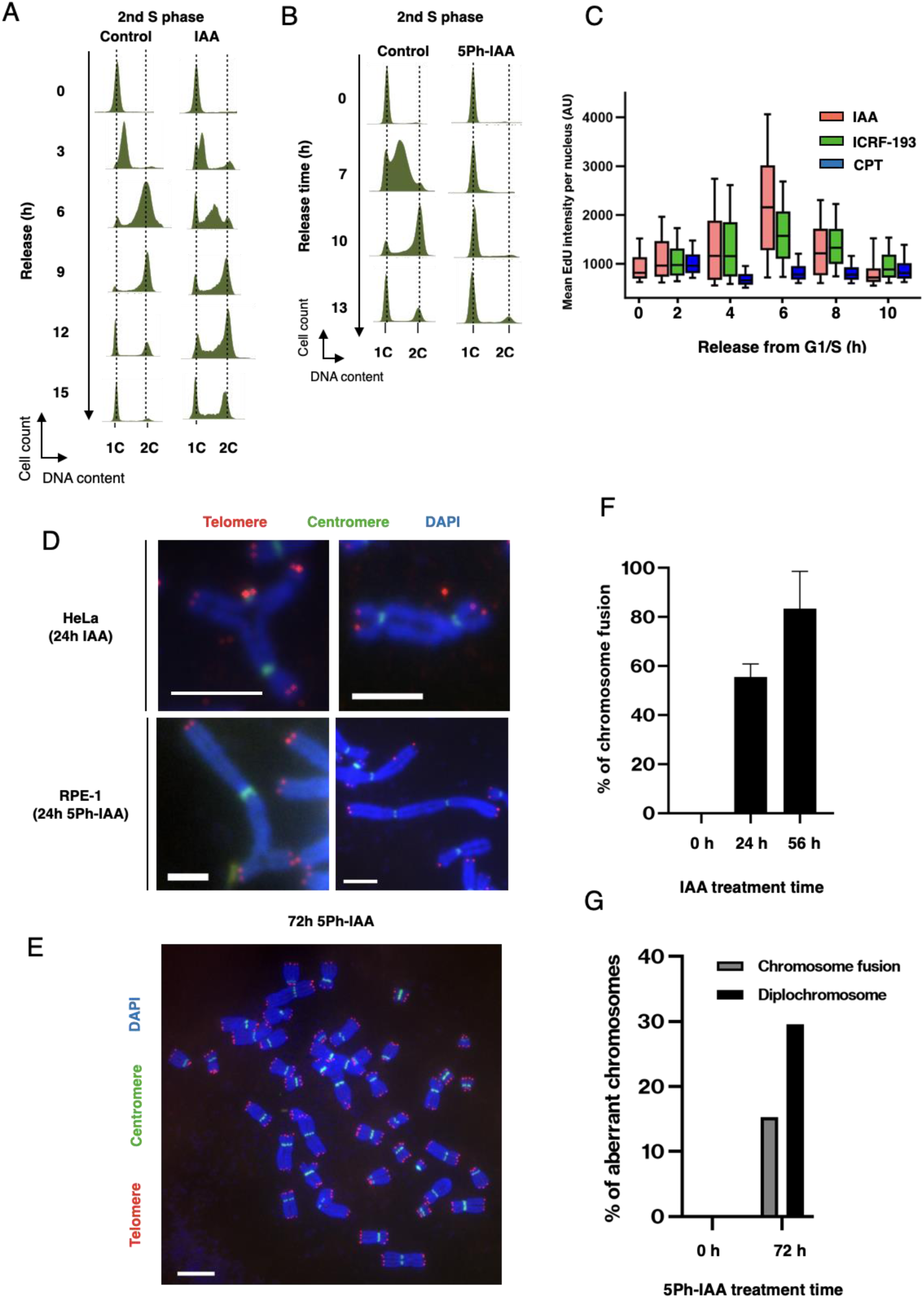
Replication perturbation and chromosome rearrangements in the absence of SMC5/6 are analogous to Top2A inactivation. (A) Flow cytometric analysis of SMC6-degron HeLa cells. The second round of replication progression from the G1/S boundary after the SMC6 depletion is shown. (B) Flow cytometric analysis of SMC6-degron RPE-1 cells. Progression through the second round of S phase after the SMC6 depletion is shown. (C) Replication of the first round of S phase in cells inhibited for topoisomerase activity. Cells were treated with IAA (red), 100 nM ICRF-193 (green), or 10 nM camptothecin (CPT) (blue) after the G1/S release. At least 1,678 cells were analyzed for each experiment. (D, E) Chromosomal rearrangements found in SMC6-depleted HeLa and RPE-1 cells. The positions of centromeres (green) and telomeres (red) were labelled by FISH in spread metaphase chromosomes. (D) Both SMC6-depleted HeLa and RPE-1 cells show chromosome fusions resulting in radial-shaped chromosomes (shown in left panels) and dicentric chromosomes (shown in right panels). (E) In RPE-1 cells, diplochromosomes were induced after SMC6 depletion. Scale bar, 10 μm. A similar procedure was used for cell collection and FISH labeling to detect chromosome fusions and diplochromosomes. (F) Bar charts summarizing the frequency of chromosome fusions in SMC6-depleted HeLa cells. At least 30 mitotic cells were analyzed in three independent experiments. (G) Bar charts summarizing the frequency of chromosome fusions and diplochromosomes in SMC6-depleted RPE-1 cells. A total of 98 mitotic cells were analyzed.

**Figure EV5.**
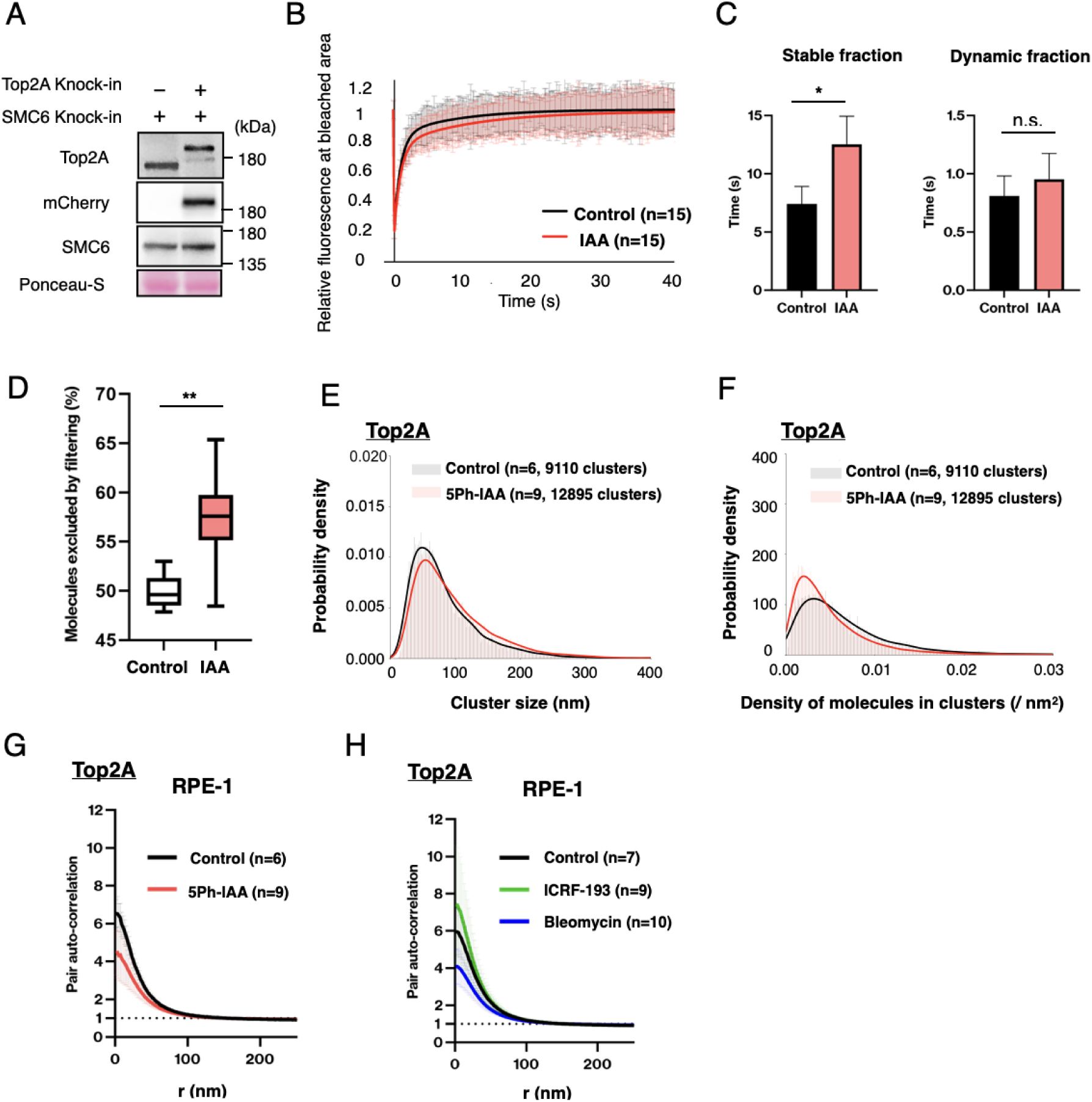
Dynamics of Top2A in RPE1 and HeLa cells. (A) Immunoblots validating endogenously mCherry labeled Top2A in SMC6-degron HeLa cells. (B, C) (B) Dynamically bound-fraction of Top2A in the nucleoplasm was slightly but repeatedly reduced by SMC6 depletion (red) from the control level (black). (C) Bar charts show the half-residence time of stably- and dynamically-bound fraction of Top2A fraction from experiments in B. Error bars represent SD of data from three independent experiments. n.s. indicates not significant. * indicates *P* < 0.05; unpaired t-test. (D-F) Quantitative analyses of Top2A cluster distributions in RPE-1 cells. Segmentation analysis identified ∼1×10^3^ Top2A clusters per cell in control and 5Ph-IAA treated RPE-1 cells in late S phase. The effects of SMC6 depletion in RPE-1 cells on the proportion of molecules excluded from clusters (D), the cluster size (E), and the cluster density (F) were shown. Two-tailed Wilcoxon-Mann-Whitney tests indicate differences in the excluded fraction (*P* < 0.01 shown as **), the size (*P* < 0.0001), and the density (*P* < 0.0001) between control and IAA. (G, H) Quantitative analysis of Top2A molecular assemblies in RPE-1 cells. The densities of Top2A assemblies in late S phase are analyzed in cells with SMC6 depletion (5Ph-IAA, red), ICRF-193 treatment (green), bleomycin treatment (blue) and control (black). These effects were similarly seen in HeLa cells (see Figure 3F, 3G).

**Figure EV6.**
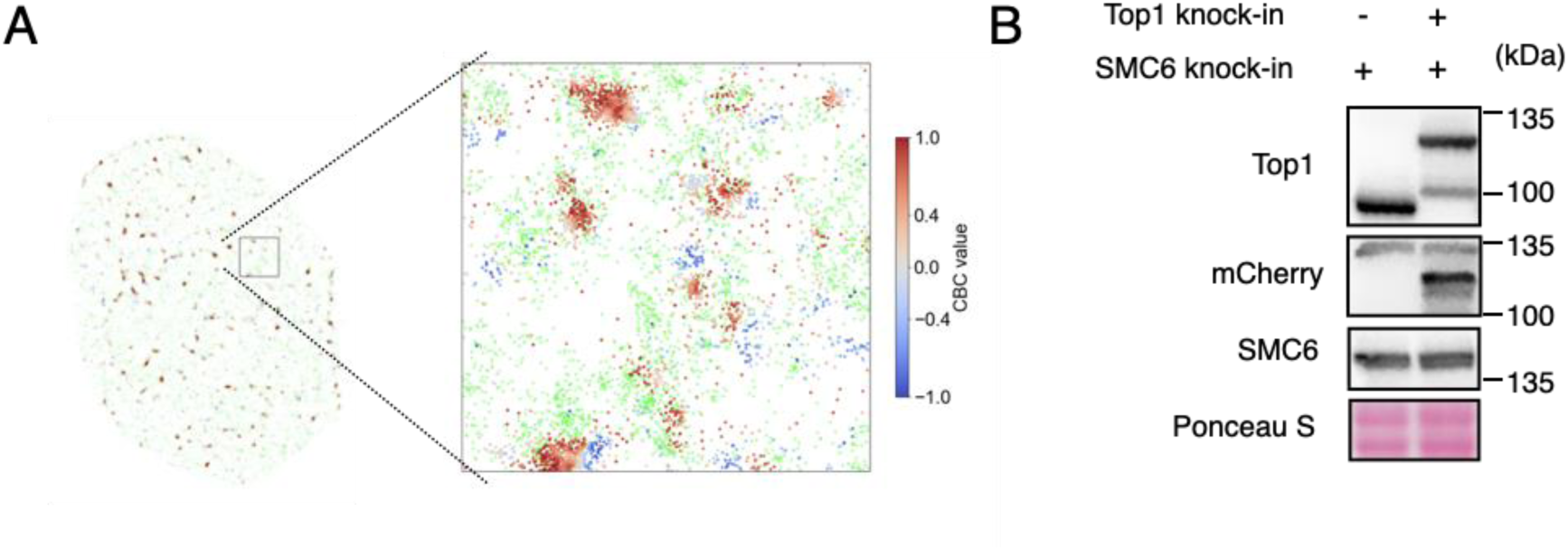
Top2A has a primary role in releasing negative supercoils associated with replication. (A) Representative color-coded density map of CBC values showing the correlated distribution of Top2A molecules to PCNA molecules in late S phase nuclei. Enlarged views show correlated distribution (red), random distribution (gray), and anti-correlated distribution (blue) with color-coded density bar are shown. Note that a fraction of Top2A is randomly excluded to examine identical number of signals with PCNA in the CBC analysis (green). (B) Immunoblots verifying endogenously mCherry-labelled Top1 in SMC6-degron HeLa cells.

**Figure EV7.**
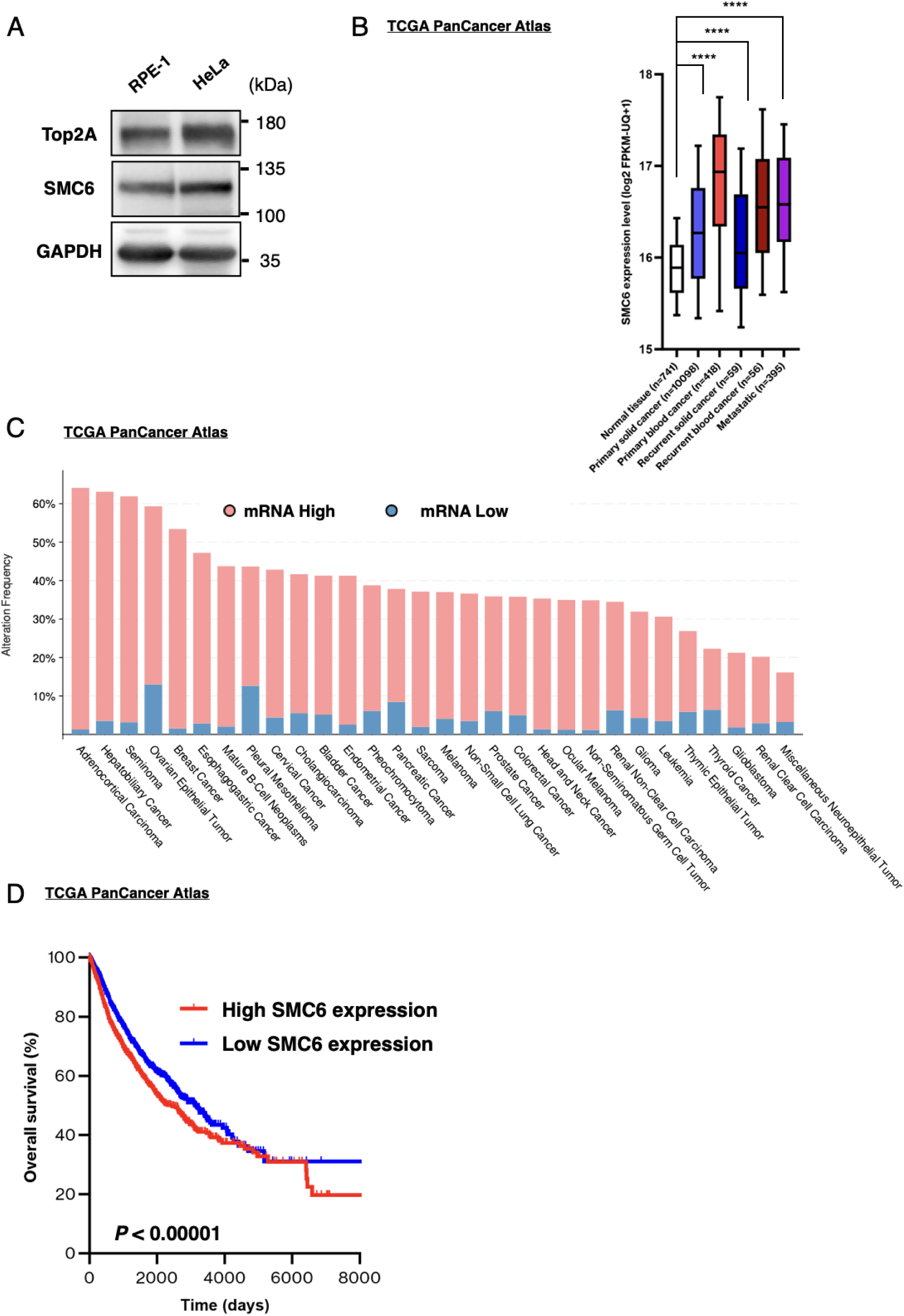
High expression of SMC5/6 and its prognostic relevance in cancer patients. (A) Immunoblots show equivalent protein levels of Top2A and SMC6 between RPE-1 and HeLa asynchronous cells. (B) A variety of cancers with or without recurrence or metastasis show a high expression level of SMC6 compared with normal tissues. **** indicates P < 0.0001; unpaired t-test. While, normal tissue; light blue, primary solid cancer; light red, primary blood cancer; blue, recurrent solid cancer; red, recurrent bold cancer; purple, metastatic cancer. mRNA expression (RNA Seq V2 RSEM) is calculated as z-score relative to diploid samples. ****, *P* < 0.0001; two-tailed Wilcoxon-Mann-Whitney tests. (C) Frequency of a high expression levels of each subunit of SMC5/6 across human cancers according to TCGA PanCancer Atlas studies. Red bar, high SMC6 expression; blue bar, low SMC6 expression. mRNA expression (RNA Seq V2 RSEM) is calculated as z-score relative to diploid samples (Figure is obtained from cBioPortal (Cerami *et al*, 2012)). (D) Kaplan-Meier curves showing overall survival of patients with primary solid and blood cancers expressing high or low levels of SMC6. Red line, patient group with high expression of SMC6; blue line, patient group with low expression of SMC6. Log-rank test indicates difference in overall survival (*P* < 0.00001).

